# Pioneer transcription factors are associated with the modulation of DNA methylation patterns across cancers

**DOI:** 10.1101/2021.05.10.443359

**Authors:** Roza Berhanu Lemma, Thomas Fleischer, Emily Martinsen, Marit Ledsaak, Vessela Kristensen, Ragnhild Eskeland, Odd Stokke Gabrielsen, Anthony Mathelier

## Abstract

Methylation of cytosines on DNA is a prominent modification associated with gene expression regulation. Aberrant DNA methylation patterns have recurrently been linked to dysregulation of the regulatory program in cancer cells. To shed light on the underlying molecular mechanism driving this process, we hypothesized that aberrant methylation patterns could be controlled by the binding of specific transcription factors (TFs) across cancer types. By combining DNA methylation arrays and gene expression data with TF binding sites (TFBSs), we explored the interplay between TF binding and DNA methylation in 19 cancer types. We performed emQTL (expression-methylation quantitative trait loci) analyses independently in each cancer type and identified 13 TFs whose expression levels are correlated with local DNA methylation patterns around their binding sites in at least 2 cancer types. The 13 TFs are mainly associated with local demethylation and are enriched for pioneer function, suggesting a specific role for these TFs in modulating chromatin structure and transcription in cancer patients. Furthermore, we confirmed that *de novo* methylation is precluded across cancers at CpGs lying in genomic regions enriched for TF-binding signatures associated with SP1, CTCF, NRF1, GABPA, KLF9, and/or YY1. The modulation of DNA methylation associated with TF binding was observed at cis-regulatory regions controlling immune- and cancer-associated pathways, corroborating that the emQTL signals were derived from both cancer and tumor-infiltrating cells. As a case example, we experimentally confirmed that FOXA1 knock-down is associated with higher methylation in regions bound by FOXA1 in breast cancer MCF-7 cells. Finally, we reported physical interactions between FOXA1 with TET1 and TET2 both in an *in vitro* setup and *in vivo* at physiological levels in MCF-7 cells, adding further support for FOXA1 attracting TET1 and TET2 to induce local demethylation in cancer cells.

## Introduction

Chromatin and DNA modifications act as molecular stamps associated with active and inactive regulatory status of corresponding genomic regions, which are crucial for proper homeostasis and development [1,2]. Among the various possible DNA modifications [3], the addition of a methyl group to the 5^th^ carbon of cytosine leads to the 5-methylcytosine (5mC) mark. The 5mC mark (hereafter referred to as DNA methylation) is usually associated with the transcriptional silencing of cis-regulatory elements such as promoters or enhancers [4,5]. As aberrant DNA methylation patterns are linked to various diseases such as cancers [6,7], it is critical to understand the underlying molecular mechanisms driving this process.

Covalent DNA methylation at cytosines (mainly in the CpG context) is acquired by the addition of 5-methylcytosine catalyzed by the DNA methyltransferase (DNMT) enzymes. DNA demethylation is carried out by the Ten-Eleven Translocation (TET) proteins in successive hydroxylation reactions resulting in 5mC derivatives [3], which are removed by thymine DNA glycosylase through the base excision repair pathway (reviewed in [8]). As DNMTs and TETs bind DNA in a limited sequence-specific manner, their recruitment to specific genomic regions has been reported to be driven by interactions with transcription factors [9–12].

Transcription factors (TFs) are proteins that recognize and bind cis-regulatory regions (promoters and enhancers) at their TF binding sites (TFBSs) through sequence-specific TF-DNA interactions to regulate transcription [13]. Through their binding at cis-regulatory regions, most TFs recruit co-factors to activate or repress the transcription of target genes [13,14]. While most of the TFs engage with open chromatin regions at their TFBSs, a specific class of TFs, the pioneer TFs, have the ability to engage with nucleosome-bound chromatin independent of other factors. Pioneer TFs are believed to be the first factors to engage with target chromatin regions and associate with compact chromatin to facilitate the binding of other additional factors and local epigenetic modifications [15–17]. For instance, changes across myeloid cell fate transitions are marked with the priming of inaccessible enhancers by pioneer TFs, which leads to locally increased chromatin accessibility and DNA methylation loss [18].

Several TFs have been reported to physically interact with DNMTs and/or TETs and are therefore likely to recruit these enzymes to specific genomic regions. The leukemogenic PML-RAR fusion protein has been shown to recruit DNMTs while RUNX1 recruits the DNA demethylation machinery [7,19,20]. Using co-immunoprecipitation in HEK293T cells and endogenous IP in LNCaP cells, FOXA1 was found to physically interact with TET1 and promote the co-occupancy of TET1 in FOXA1 occupied regions [21].

To investigate the association between TF binding and DNA demethylation at large scale, Suzuki *et al*. developed a screening system combined with TF binding motif enrichment at differentially methylated regions after ectopic expression of selected TFs. This strategy identified a set of developmental (cell fate determining) TFs that were associated with binding site-directed DNA demethylation [6]. Another high-throughput screening strategy investigated the interplay between TF binding and DNA methylation for hundreds of TFs [22]. The strategy relies on the integration of a sequence backbone with known methylation status but containing diverse TF binding motifs followed by bisulfite sequencing of PCR amplicons. The study revealed pioneer TFs that can induce local DNA demethylation and pioneer TFs whose binding have a protective effect against *de novo* DNA methylation [22]. Using a computational approach, the ELMER (Enhancer Linking by Methylation/Expression Relationships) tool allowed for the large-scale identification of transcriptional enhancers and their target genes based on DNA methylation data (at enhancers) and gene expression [23]. Motif enrichment analysis at the enhancers predicted pan-cancer by ELMER inferred TFs that could act as upstream regulators of DNA methylation patterns at these enhancers [23]. Despite continuous efforts to unravel the molecular mechanism by which DNA methylation is regulated, the current understanding of how DNA methylation is regulated and its interplay with TF-binding in cancer patients is limited [6,7,18]. We hypothesized that a pan-cancer and genome-wide investigation of the interplay between TF binding and resulting local DNA methylation patterns in cancer genomes could reveal key regulatory processes that are critical for an improved molecular understanding of cancers.

In this study, we designed a computational approach to identify CpGs with DNA methylation level correlated with the expression level of 231 TFs. We further assessed the enrichment of these CpGs around TFBSs for the corresponding TFs. This TF binding-centric expression-methylation quantitative trait loci (emQTL) methodology was applied to 19 cancer types from The Cancer Genome Atlas (TCGA) to predict TFs associated with DNA methylation patterns (emTFs, expression-methylation TFs). The analyses revealed 13 emTFs (33 TF-cancer type pairs) for which an enrichment for correlated CpGs around their TFBSs was observed in at least 2 cancer types, providing evidence for their potential role in DNA methylation patterns in cancer patients. The pioneer function of these 13 emTFs, which we found predominantly associated with DNA demethylation, has been demonstrated by previous studies. Furthermore, we confirmed the presence of TF-binding signatures that are discriminative between regulatory regions associated with varying DNA methylation across patients and regions where *de novo* DNA methylation is precluded. From the list of 13 emTFs, we experimentally investigated the role of FOXA1 in DNA demethylation in breast cancer. We observed that FOXA1 knockdown led to an increase of DNA methylation at some regions bound by FOXA1 in MCF-7 cells. We further reported physical interactions between FOXA1 and both TET1 and TET2 at physiological levels in MCF-7 cells as well as using *in vitro* GST-pulldown assays.

## Results

### Prediction of transcription factors associated with DNA methylation patterns around their binding sites across cancer types

We aimed to unravel the interplay between TF binding to the DNA and local DNA (de-)methylation. We hypothesized that the binding of specific TFs to their TFBSs would be correlated with local DNA (de-)methylation if these factors were associated with DNA modifications. By combining DNA methylation (from Illumina 450k arrays) and gene expression data from 19 cancer type cohorts from TCGA [24] (Table S1) with high-quality direct TF-DNA interactions (i.e. TFBSs) from the UniBind database [25], we assessed the correlation between DNA methylation and TF-binding using TF expression as a surrogate for TF binding potential at their TFBSs. Altogether, we evaluated the expression of 231 TFs with DNA methylation at CpGs in cancer cohorts of 59 to 703 patients (Table S1). Specifically, we performed expression-methylation quantitative trait loci (emQTL) analyses by computing Spearman correlation coefficients between the expression of the 231 TFs and methylation level at 376,997 CpGs located close to TFBSs in each cancer type independently (see Materials and Methods for details and Table S2 for the number of CpGs close to TFBSs for each TF). This emQTL computation followed our previously published methodology associating CpGs with gene expression [26] but was restricted to TFs and CpGs surrounding their binding sites. Note that for each TF, we considered all 376,997 CpGs in the emQTL analysis.

For each TF we examined the proportion of the CpGs close to its TFBS that were in emQTL with the TF itself; the percentages varied significantly between TFs and across cancer types (Figure 1A). In some cancer types, several TFs were associated with high percentages of correlated CpGs (e.g. in breast cancer, BRCA, and brain lower grade glioma, LGG) while small proportions of CpGs were observed for all TFs in other cancer types (e.g. in glioblastoma multiforme, GBM, and acute myeloid leukemia, LAML). We examined whether this variability could be explained by the lack of statistical power in the emQTL analyses for the cohorts with a lower number of samples. Indeed, we observed a significant correlation between the number of samples in a cohort and the median number of correlated CpG percentages (Figure S1B). We speculate that the identification of TFs that could be associated with local DNA methylation patterns around their TFBSs is precluded in cohorts with smaller sample sizes.

**Figure 1.**
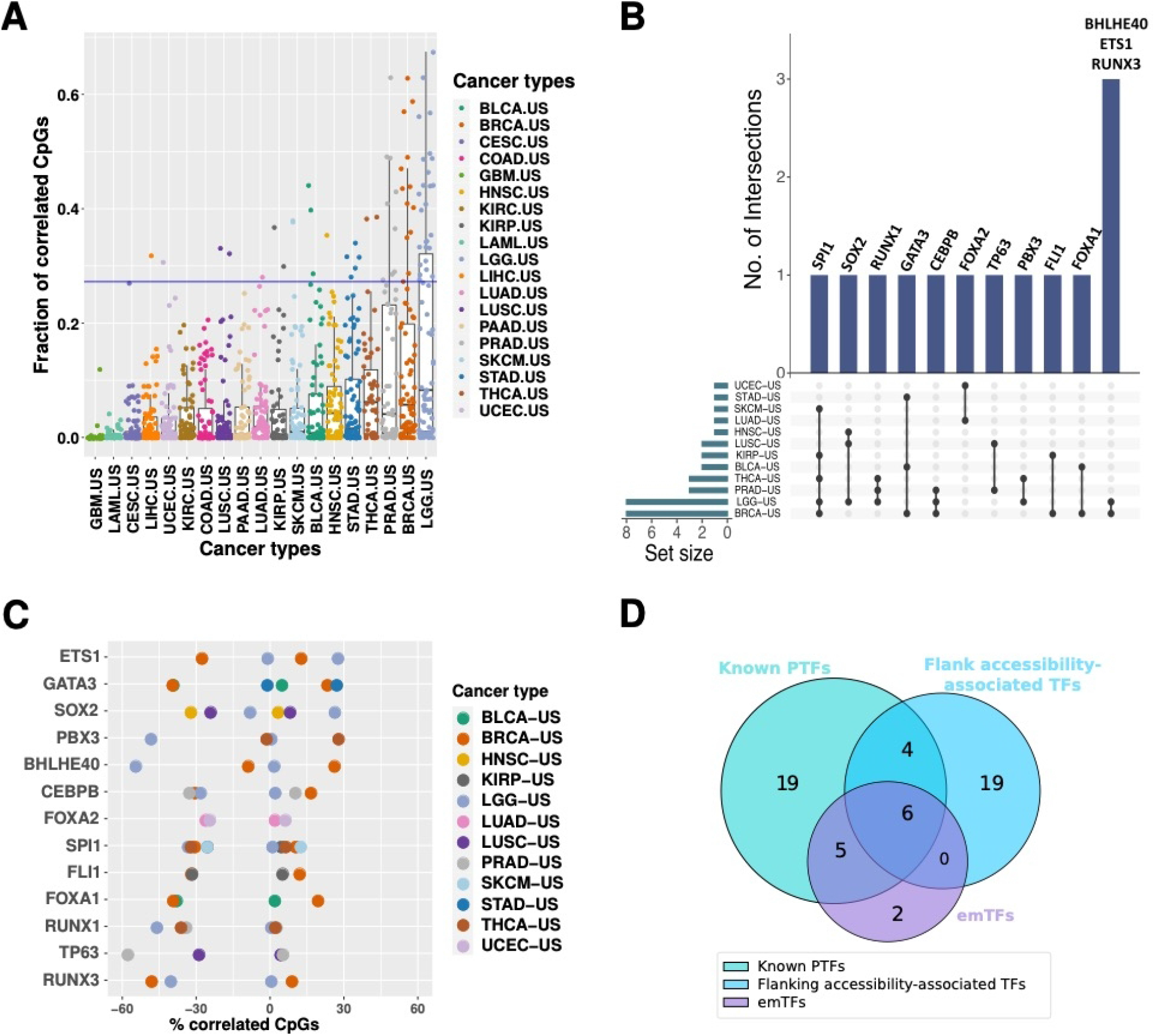
Identification and analysis of emTFs. **A**. Box plot depicting the fraction of CpGs close to TFBSs for each TF (each point corresponds to a TF in a given cohort (columns)) with DNA methylation level correlated with the TF expression. The horizontal blue line represents the 95th percentile of the distribution of all fractions (see Figure S1 for the distribution). **B**. Upset plot representing the emTF predictions across the cancer types. Each row represents a cancer type with points providing information about the intersection of the TFs predicted in the different cancer types. The bars at the top indicate the number of intersecting TFs (annotated above each bar) in each combination of cancer types (indicated by the points). The set size below the horizontal bars depicts the number of TFs predicted in each cohort. **C**. For each emTF (rows), the plot provides the percentages of correlated CpGs (emCpGs; x-axis) close to their TFBSs predicted in each cancer type (see legend). emCpGs are further decomposed into the ones exhibiting a positive or negative correlation between their methylation level and the TF expression. The proportion of positively correlated emCpGs is provided as a positive value (x-axis) while the proportion of negatively correlated emCpGs is provided with negative values (x-axis). See Table S3 for the detailed numbers **D**. Venn diagram of the intersection between the predicted emTFs (n=13), pioneer TFs from the literature (known PTFs; n=34), and flanking accessibility-associated TFs from [40] (n=29).

To focus on the TFs for which the binding is most likely to have a local effect on DNA methylation, we considered the TFs associated with the highest percentages of correlated CpGs that were especially enriched close to their TFBSs. Specifically, we extracted the top 5% of CpG percentages from the distribution obtained for all TF-cancer pairs (Figure S1C). In addition, we filtered out TFs that did not show a specific enrichment of CpGs in emQTL close to their TFBSs. The filtering was achieved by assessing the enrichment for correlated CpGs around the TF’s TFBSs using Mann-Whitney U tests; we retained TFs with p-values < 0.01 (Materials and Methods; Table S3). This strategy revealed 37 TFs in 12 cancer types (Figure S1D). We observed consistent association with local DNA methylation patterns in at least 2 cancer types for 13 TFs (Figure 1B). Even though the 13 TFs were associated with an enrichment of correlated CpGs close to their binding sites, the corresponding CpGs identified in each cancer type vary (Figures S2-S3). Hereafter, we refer to these 13 TFs as emTFs (expression-methylation TFs) and to the correlated CpGs close to their TFBSs as emCpGs (expression-methylation CpGs).

Cytosines represented in the Illumina 450K methylation array are not distributed evenly throughout the genome but mainly localised in proximal promoters and gene bodies [27]. Similarly, the TFBSs from UniBind that were considered in this study are also predominantly found at proximal promoters [28]. We assessed the genomic distribution of emCpGs and compared it to the complete set of 376,997 CpGs considered (and located close to TFBSs). Across cancer types, we observed a smaller proportion of emCpGs at proximal promoters than observed with the complete set of CpGs, while emCpGs were more frequently found at intronic and intergenic regions (Figure S4). This observation suggests that emCpGs are more predominantly detected at enhancer than promoter regions.

### emTFs are mainly associated with demethylation and are enriched for pioneer function

We sought to provide molecular mechanistic insights underlying the interplay between emTF binding and local DNA methylation modulation around their TFBSs. We first investigated the nature of the correlations (positive versus negative correlations) between emTFs’ expression and DNA methylation at emCpGs. Across cancer types, the expression of the emTFs was mainly negatively correlated with the level of methylation of the associated emCpGs (Figure 1C; Figure S5). The proportion of negatively correlated emCpGs ranged from ∼1% to ∼58% per TF-cohort (mean=30.7%; median=32.1%), while the proportion of positively correlated emCpGs ranged from ∼0.3% to ∼28% (mean=9.6%; median=6%). These results indicate that, in most cases, higher emTF expression is associated with lower CpG methylation around their TFBSs, suggesting local DNA demethylation through TF binding.

Higher level of DNA methylation is usually associated with silenced and inaccessible cis-regulatory regions [4,29]. We speculated that the emTFs would engage with these regions of methylated and closed chromatin to trigger demethylation and chromatin accessibility. As pioneer TFs have the capacity to engage with closed chromatin, we examined if the 13 identified emTFs were enriched for such pioneer function. We collected a list of pioneer TFs by reviewing the literature (Table S4) [30–39] and found that the emTFs were enriched in the list of pioneer TFs (11 out of the 13 emTFs; Fisher test p-value < 9.4e^-31^; Figure 1D).

Next, we aimed to provide complementary evidence for emTFs to engage with closed chromatin and reshape the chromatin landscape in cancer patients. A recent study reported the chromatin accessibility landscape of human cancers using ATAC-seq [40]. This work predicted 55 TFs (29 of which were among the 231 TFs investigated in this study) for which the binding is associated with increased chromatin accessibility in the regions flanking their TFBSs, providing evidence for their pioneer function [40,41]. We found that the emTFs were enriched in the list of flanking accessibility-associated TFs reported from cancer samples in [40] (6 out of the 13 emTFs: CEBPB, GATA3, FOXA1, RUNX1, RUNX3, and TP63; Fisher test p-value < 3.9e^-15^; Figure 1D; Table S5).

Furthermore, the ATAC-seq study observed that the increased chromatin accessibility was accompanied by local DNA demethylation [40]. For each cancer type, we considered the emCpGs lying in open chromatin regions and computed spearman correlations between their level of methylation and the level of openness of the regions that contain them (Materials and Methods). As expected, we recapitulated the results previously observed [40] with consistent negative correlations between chromatin accessibility and DNA methylation at emCpGs (Figure S6). Although the number of matching patient IDs for the other cancer types investigated is too small, we still observed similar correlation trends.

Taken together, these results provide complementary supporting evidence for the enrichment of emTFs with pioneer function to promote chromatin accessibility and demethylation in a binding site-directed fashion in cancer patients.

### *De novo* methylation-protected CpGs and CpGs associated with emTFs harbour distinct TF-binding signatures

In the previous sections, we revealed that regions around emTF binding sites harboured significant proportions of emCpGs. Nevertheless, not all CpGs proximal to the corresponding TFBSs exhibited DNA methylation levels correlating with the emTFs’ expression across patients. We investigated whether distinct TF binding patterns could discriminate between these two sets of CpGs (correlated/emCpGs versus uncorrelated for each emTF in each cancer type). For each emTF-cancer type pair, we looked for the differential enrichment of TFBSs for 231 TFs using the UniBind enrichment tool [28], when considering regions surrounding emCpGs versus non-correlated CpGs and vice-versa (Materials and Methods).

We consistently observe that regions of ± 200 bp surrounding emCpGs for a given emTF are differentially enriched for binding sites bound by that particular emTF (Figures 2A and Supplementary Data 1). It is important to note that both emCpGs and non-correlated CpGs are close to TFBSs for the given emTF and the regions analyzed did not exhibit distinct %GC content (Figures 2C, S7A-S17A). Hence, the differential enrichment analysis highlights that regions flanking emCpGs contain significantly more TFBSs for the emTF than regions flanking non-correlated CpGs, without an overall nucleotide composition difference. Figure 2A depicts a representative example using flanking regions of CpGs close to FOXA1 TFBSs with emCpGs and non-correlated CpGs identified in the BRCA-US cohort. Note the combined enrichment for FOXA1, ESR1, and GATA3 TFs close to the emCpGs; these 3 TFs have already been associated with DNA methylation patterns in estrogen receptor positive breast cancers [26].

**Figure 2.**
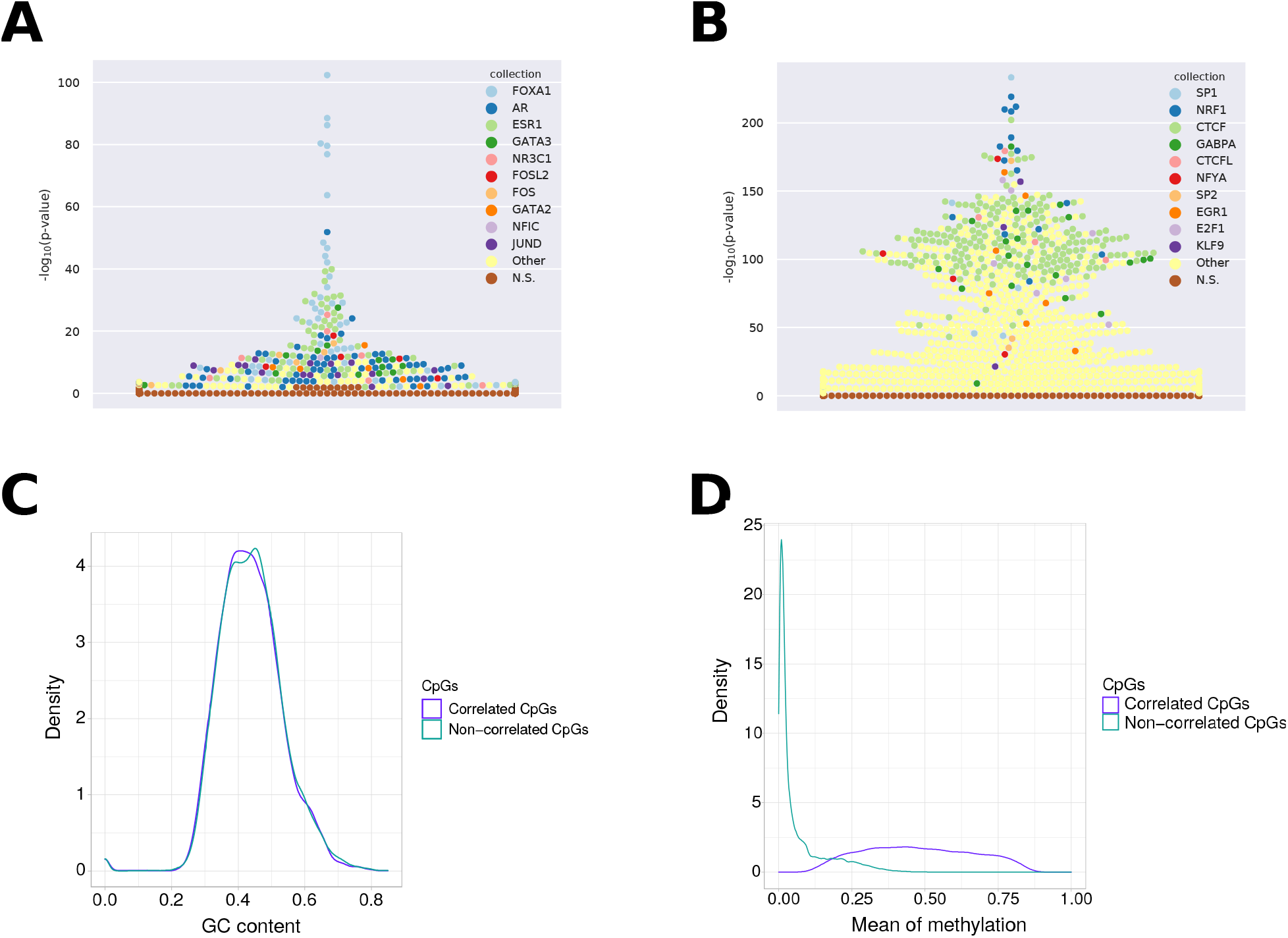
TF binding signatures at FOXA1-associated emCpGs versus *de novo* methylation-protected CpGs in breast cancer. **A**. Beeswarm plot depicting TFBS sets enrichment (y-axis) specific to regions surrounding emCpGs associated with FOXA1. Each point corresponds to a TFBS dataset in UniBind (one color per TF, see legend). **B**. Beeswarm plot depicting TFBS sets enrichment (y-axis) specific to regions surrounding non-correlated CpGs, which are close to FOXA1 TFBSs but whose DNA methylation levels do not correlate with FOXA1 expression in breast cancer samples. **C**. Density distribution (y-axis) of GC contents (x-axis) at regions surrounding FOXA1 emCpGs (purple) and non-correlated CpGs (green). **D**. Density distribution (y-axis) of mean methylation levels (x-axis) across breast cancer samples for FOXA1 emCpGs (purple) and non-correlated CpGs (green).

The analyses of regions surrounding non-correlated CpGs consistently revealed the differential enrichment for TFBSs associated with the TFs CTCF, YY1, NRF1, GABPA, KLF9, and SP1 (Figures 2B and Supplementary Data 2). The enrichment of these TFs is in agreement with previous studies that identified the binding of SP1, CTCF, NRF1, and YY1 to prevent *de novo* methylation [42–45]. The protective effect of these TFs against *de novo* methylation is in line with the constant hypomethylation of the non-correlated CpGs observed across emTFs and cancer cohorts (Figures 2D and S7B-S17B).

Altogether, these results support the existence of two distinct TF binding signatures that discriminate emCpGs associated with emTFs from other CpGs close to the TFBSs of emTFs. While the emCpGs harbour enriched binding sites for their specific emTFs, the non-correlated CpGs shared a binding signatures for SP1, CTCF, NRF1, GABPA, KLF9, and YY1 providing a protective effect against *de novo* methylation across cancer types.

### emCpGs are predicted to regulate genes involved in immune response, cell fate determination, and cancer pathways

With multiple lines of evidence supporting the pioneer function of the emTFs, we hypothesized that they might be involved in the activation of specific genes via demethylation of emCpGs in cis-regulatory regions. As the methylation and expression data from TCGA were derived from bulk tumours, the samples are a combination of cancer cells and cells from the tumour microenvironment. Hence, some emTFs might be acting upon cancer cells while others would be active in cells from the microenvironment. We investigated the association between the observed emQTL signals and tumour purity of the TCGA samples. By comparing the level of expression of the emTFs with the tumour purity estimate of the samples in the cancer cohorts, we observed a positive correlation for about half of the emTFs (Materials and Methods; Figures S7C-17C). emTFs BHLHE40, ETS1, FOXA1, FOXA2, GATA3, PBX3, TP63, and SOX2 lie in this category across several cancer types (Figures S7C-17C). The positive correlation points to the emQTL signal being mostly driven by cancer cells in the associated cohorts. On the contrary, the expression of some emTFs in specific cancer types was negatively correlated with tumour purity (Figure S7C-17C). emTFs CEBPB, ETS1, FLI1, BHLHE40, TP63, GATA3, PBX3, RUNX1, RUNX3, and SPI1 lie in this category across several cancer types (Figures S7C-17C). The negative association with tumour purity indicates that these emTFs might be acting in cells from the microenvironment in the corresponding cancer types.

To assess the functional relevance of the identified emCpGs in these different cellular contexts, we estimated the enrichment for biological processes and pathways in the list of genes linked to emCpGs for each pair of emTF-cancer types. We linked emCpGs to genes using gene-specific regulatory elements defined by the STITCHIT algorithm, which relies on an integrative analysis of epigenetic and transcriptomic data [46]. This method allows to assign emCpGs lying in distal *cis*-regulatory elements to their potential target genes. When emCpGs did not lie within STITCHIT regulatory elements, we assigned them to the closest gene (Materials and Methods). The emCpG-gene links were derived from multiple cell types / tissues but we aimed to focus on the most likely regulatory links in a cancer type-specific way. Specifically, we required a significant (Bonferroni adjusted p-value < 0.01) negative correlation between emCpG methylation level and target gene expression in a given cancer type to conserve an emCpG-to-gene link.

The genes linked to emCpGs associated with cancer-cell emTFs were mostly found enriched in hormone- and cancer-associated Hallmark sets of genes from the Molecular Signatures Database [48] (MSigDB; Figure 3C). For instance, emCpGs associated with FOXA1, FOXA2, and GATA3 were linked to genes enriched in estrogen receptor signaling pathways; SOX2 emCpGs enriched for genes associated with apoptosis; ETS1 emCpGs enriched for genes associated with epithelial to mesenchymal transition (Figure 3C). Moreover, we observed the recurrent enrichment for genes in Gene Ontology biological processes (GO-BP) associated with cell fate determination and development (i.e. differentiation-, development-, morphogenesis-, and growth-related terms; Figure S18A). The enrichment for these processes is in line with the biological function of pioneer TFs, which are associated with the control of cell fate and cell lineage reprogramming in normal development and cancers [15,30,49,50].

**Figure 3.**
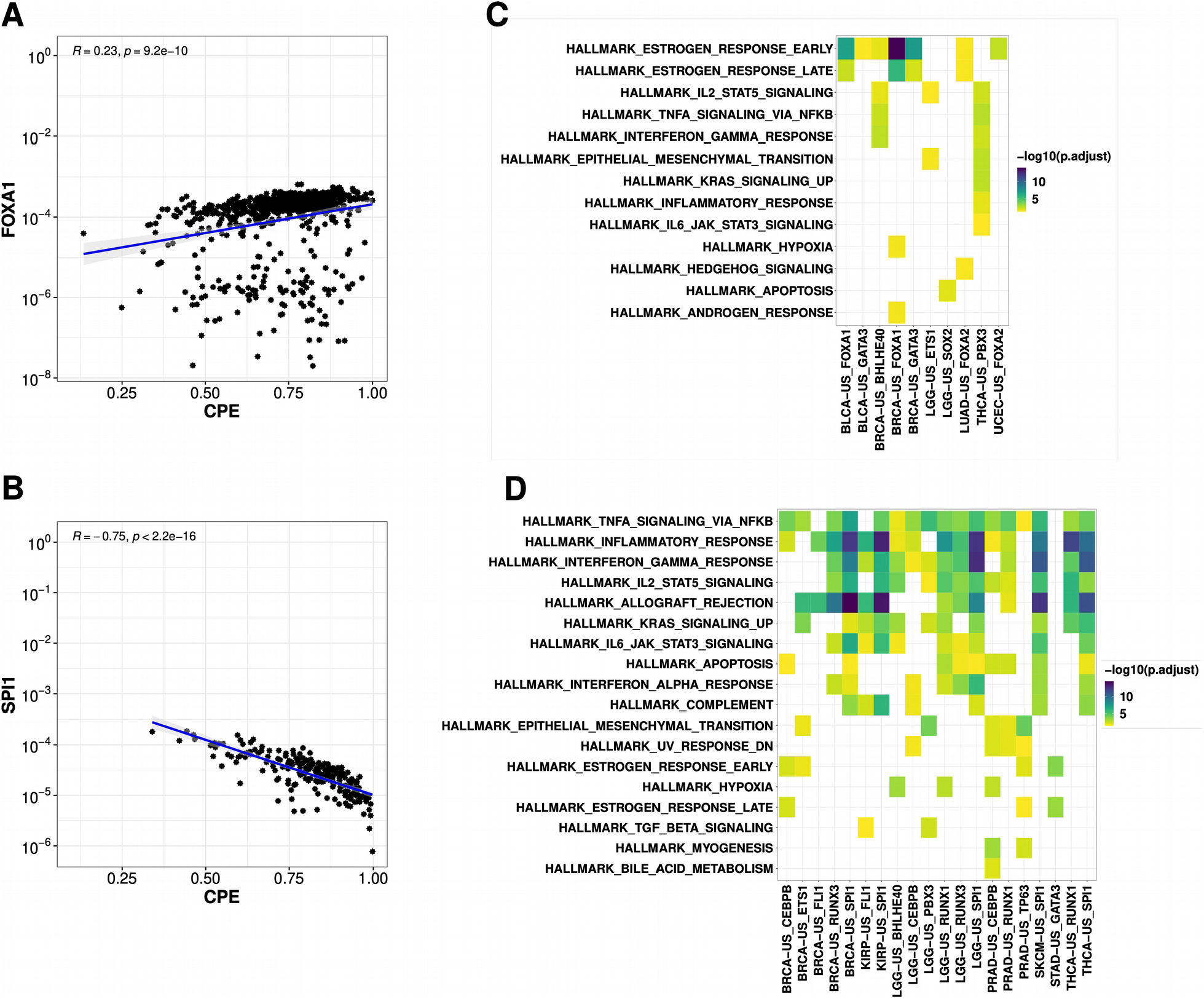
Functional evaluation of the emCpG gene targets. **A**. Pearson correlation between FOXA1 expression and tumor purity in BRCA patients **B**. Pearson correlation between SPI1 expression and tumor purity in KIRP patients. As a tumor purity variable, we used cumulative purity estimate (CPE) from BRCA and KIRP patients, respectively reported by Aran et al. [47]. The scatterplots compare the tumour purity (x-axis; cumulative purity estimate ;CPE) and expression of the TFs (y-axis). The blue lines represent the fitted Pearson linear relationship with the grey zone representing the 95% confidence interval (Pearson R coefficients and associated p-values are provided in the top-left corner). The expression of FOXA1 in breast cancer patients shows positive correlation indicating that the signals observed in the GO term and pathway enrichments are coming from the tumor cells themselves. The expression of SPI1 in kidney renal papillary cell carcinoma patients shows negative correlation indicating that the signals observed in the GO term and pathway enrichments may be coming from the tumor microenvironment. **C**. Functional enrichment analysis for genes linked to emCpGs associated with cancer cell emTFs (i.e. emTFs whose expression positively correlate with tumor purity as in A.). **D**. Functional enrichment analysis for genes linked to emCpGs associated with immune cell emTFs (i.e. emTFs whose expression negatively correlated with tumor purity as in B.). Functional enrichments in C-D were performed using the Hallmark sets from MSigDB [48].

When considering emCpGs linked to emTFs associated with cells from the tumour microenvironment, we observed the recurrent functional enrichment for immune-related terms both from MSigDB and GO-BP (Figures 3D, S18B). The functional enrichment observed suggests that the emQTL signal associated with these emTFs in the corresponding cancer cohorts is derived from tumour infiltrating lymphocytes.

Taken together, these results highlight that some emTFs are likely associated with immune cells in the tumour microenvironment while other emTFs are likely driving local demethylation of targeted cis-regulatory regions.

### Experimental assessment of the impact of FOXA1 expression on DNA methylation in MCF-7 breast cancer cells

We sought to experimentally assess the impact of the expression of an emTF on DNA methylation around its TFBSs using a cancer cell line. We selected FOXA1 and evaluated the impact of its expression in the MCF-7 breast cancer cell line. Specifically, we profiled DNA methylation in MCF-7 cells using Illumina EPIC methylation arrays under three conditions in triplicate: (1) control, (2) endogenous knock-down (KD) of FOXA1, and (3) rescue of the endogenous KD by transient ectopic expression of FOXA1-V5 (see Figure S19 for evaluation of the KD and transient rescue efficiencies using western blot). Compared to the control condition, the KD experiment assessed DNA methylation with less FOXA1 proteins, while the transient ectopic expression of FOXA1-V5 was used to try to rescue endogenous expression of FOXA1 after KD and to evaluate how it could restore the DNA methylation phenotype observed in the control condition.

We specifically evaluated the effect of FOXA1 KD on DNA methylation at genomic regions observed to be bound by FOXA1 in MCF-7 cells captured by ChIP-seq experiments (Materials and Methods). DNA methylation levels of the 83,521 CpGs within FOXA1 ChIP-seq peak regions were compared between control and KD replicates with the mCSEA tool [51] to identify differentially methylated regions (DMRs; see Material and Methods). mCSEA predicted 229 DMRs (adjusted p-value < 0.1), encompassing 431 CpGs. We observed that CpGs within the DMRs mostly exhibited higher levels of methylation after FOXA1 KD (Figure 4A). Rescuing FOXA1 expression using transient ectopic expression of FOXA1-V5 did not restore methylation at the identified DMRs after 24 hours (Figure 4A). The lack of demethylation observed after 24 hours of ectopic expression of FOXA1-V5 might be due to a slow DNA methylation process as previously observed [12]. Figures 4B-D provide case examples of DMRs after FOXA1 KD in the promoter regions of genes that have previously been associated with breast cancer: *GREB1* (growth regulation by estrogen in breast cancer 1, a regulator of hormone-dependent breast cancer growth [52]), *TFF1* (trefoil factor 1, an estrogen-regulated protein [53]), and *BRIP1* (BRCA1 Interacting Protein C-Terminal Helicase 1, which mutants participates in breast cancer development [54]).

**Figure 4.**
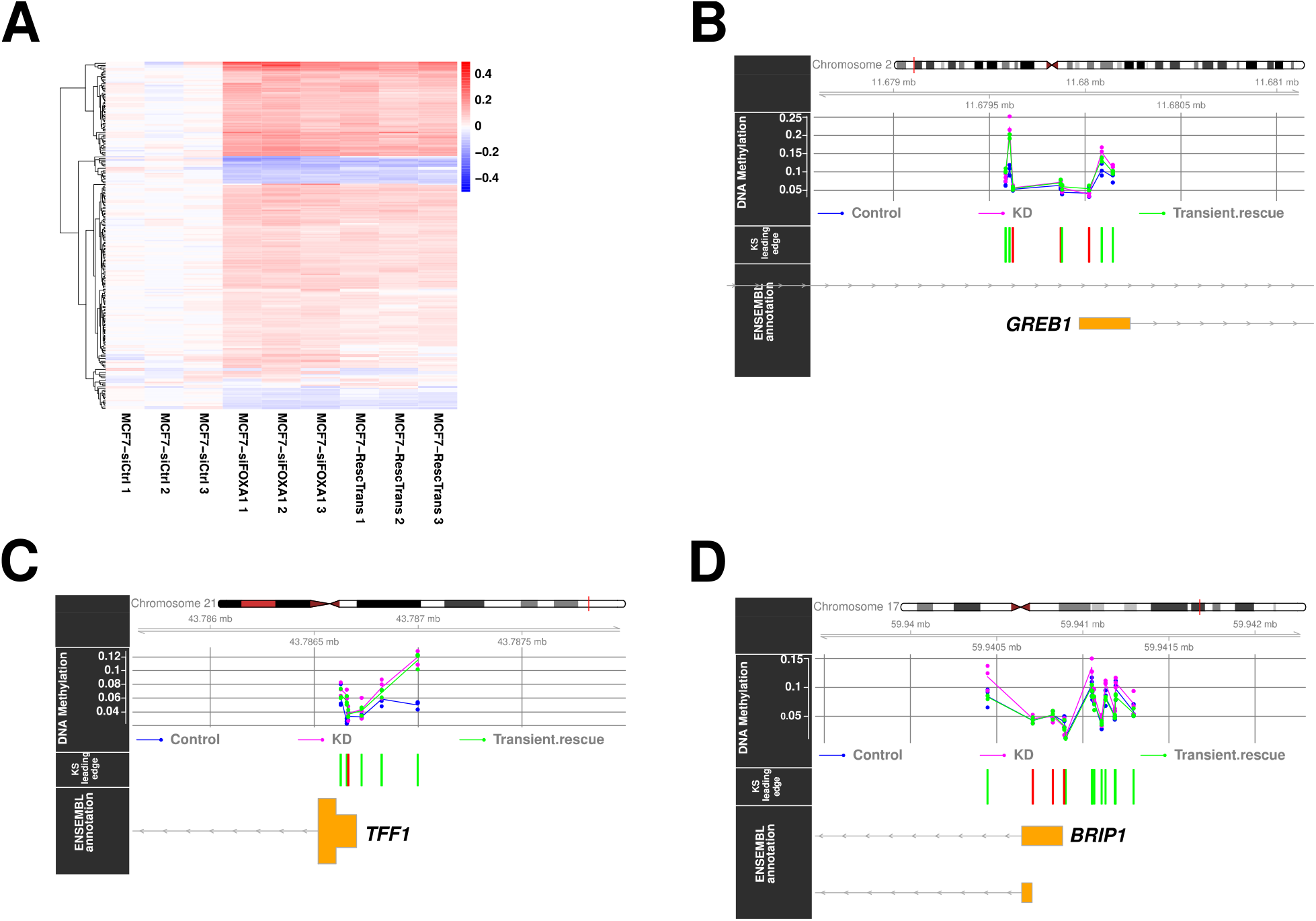
FOXA1 KD in MCF-7 cells leads to local DNA methylation increase. **A**. Heatmap depicting DNA methylation - values at 228 CpGs (rows) in DMRs (the mean -value at the control replicates are subtracted to the -value of each CpG, see Material and Methods). Blue indicates demethylation when compared to the control replicates and red indicates increased methylation. See Figure S18A for immunoblotting evaluation of the three conditions (control, KD, and rescue). **B, C, D**. Genomic context and methylation information at 3 of the 229 identified DMRs, which correspond to the promoter regions of the *GREB1* (B), *TFF1* (C), and *BRIP1* (D) genes. The upper panels provide the location in the corresponding chromosome. The second panel from the top provides the beta-values at the CpGs in the regions (blue for control samples, purple for knock-down (KD), and green for transient rescue). The third panel from the top indicates in green the significant CpGs (green) used to predict the DMR while the non-significant CpGs are depicted in red.

The experimental results outlined here confirm the association between FOXA1 expression and DNA methylation levels at genomic regions bound by FOXA1. The KD of FOXA1 increased methylation at regions bound in MCF-7 by FOXA1, supporting the link between FOXA1 binding and local demethylation.

### FOXA1 physically interacts with TET1 and TET2 at endogenous levels

The observations above suggest that FOXA1 is associated with demethylation, which can be achieved by the TET1 and/or TET2 proteins. While FOXA1 has been shown to interact with TET1 in LNCaP (lymph node carcinoma of the prostate) cell lines [21], no interaction has been reported in breast cancer cell lines with neither TET1 nor TET2, to the best of our knowledge. We aimed to assess potential protein-protein interactions between FOXA1 and TET1 and/or TET2 in MCF-7 cell lines.

We first assessed interactions for FOXA1 with TET1 and TET2 *in vitro* through GST-pulldown assays. The assays were performed using N-terminally GST fused full length human TET1 and mouse TET2 isoform 2 (mTET2) (Materials and Methods). Note that the mTET2 aligns well with the C-terminal half of the human TET2 (from residue 1388 to 2002, see Figure S20A for protein sequence alignment and Figure S20B for structural alignments of hTET2 and mTET2). Using the GST tagged TET proteins, we successfully pulled out FOXA1-V5 from COS-1 cells whole protein extract where FOXA1-V5 was transiently transfected in these cells for 24h (Figure 5A,B).

**Figure 5.**
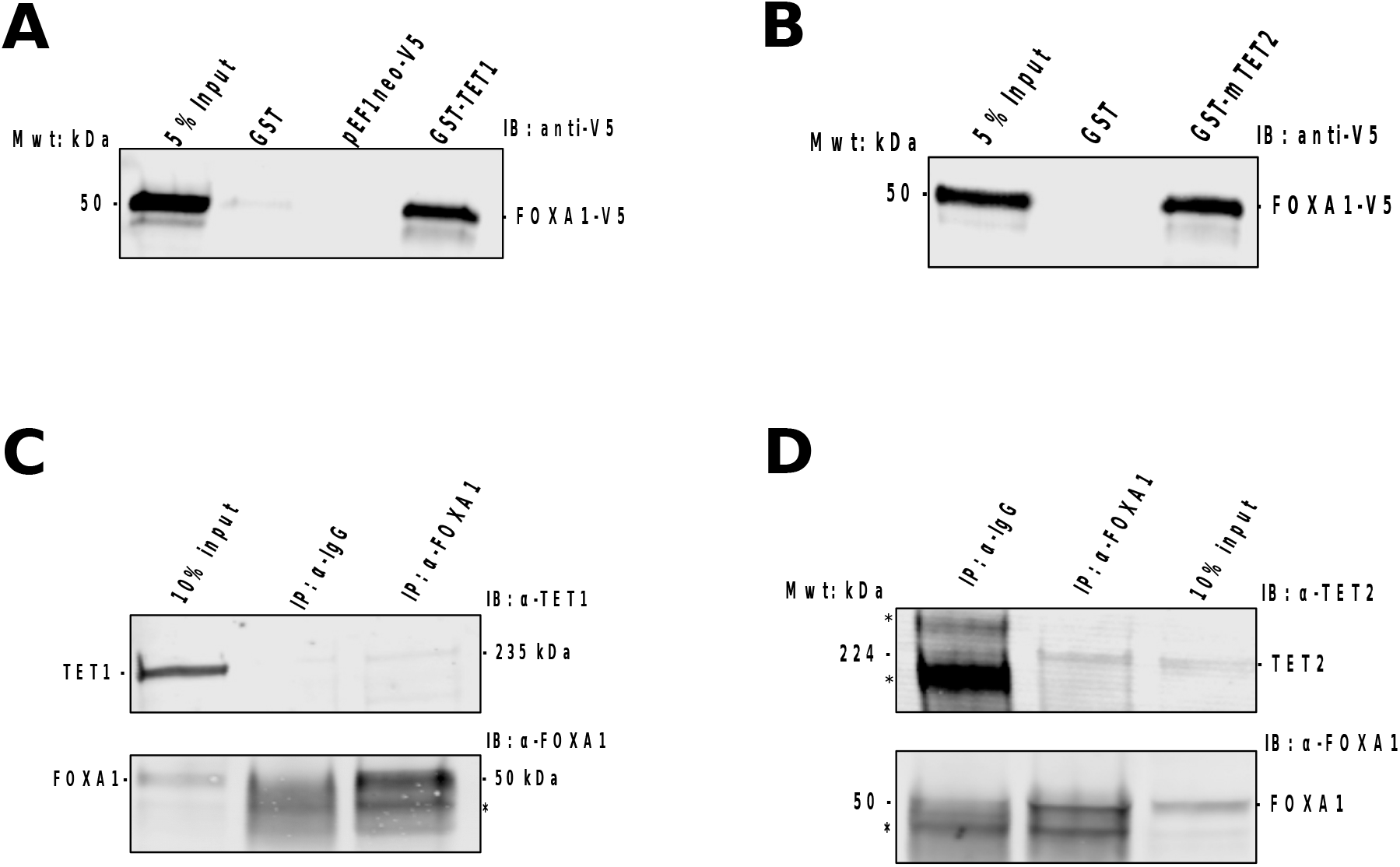
FOXA1 endogenous interaction with TET1 and TET2. We employed endogenous IP on nuclear extracts obtained from MCF-7 cells. **A**. GST-pulldown of FOXA1-V5 using GST-TET1 **B**. GST-pulldown of FOXA1-V5 using GST-mTET2 **C**. Immunoprecipitation of endogenous FOXA1 using rabbit anti-FOXA1 antibody to detect endogenous TET1 pulled together with endogenous FOXA1. **D**. Immunoprecipitation of endogenous FOXA1 using rabbit anti-FOXA1 antibody and detecting endogenous TET2 pulled together with FOXA1. Non-specific bands are marked with asterisks (*).

Next, we investigated the interactions between FOXA1 and the TETs *in vivo* in MCF-7 cells. We performed endogenous immunoprecipitations using nuclear extracts derived from MCF-7 cells (Materials and Methods). Immunoprecipitation of FOXA1 revealed an interaction with TET1 and TET2 endogenously at physiological levels in MCF-7 cells (Figure 5C,D).

Taken together, these results suggest that TET1 and TET2 interact with FOXA1 and that they are recruited by FOXA1 in MCF-7 cells for local demethylation. These interactions further support the *in silico* predictions for the importance of FOXA1 in driving local demethylation patterns in breast cancer.

## Discussion

We established a computational framework that allowed for a systematic investigation of the interplay between TF binding and DNA methylation in cancer patient samples. Through emQTL computations, we predicted 13 TFs to be associated with DNA methylation patterns around their binding sites across several cancer types. We confirmed that specific genomic regions are protected against *de novo* methylation and harbour a characteristic TF binding motif signature with enrichment of binding sites for SP1, CTCF, NRF1, GABPA, KLF9, and/or YY1. The 13 emTFs are strongly enriched for TFs with previously established pioneer function, which enables them to engage with closed chromatin and reshape the chromatin landscape. We found that some of the emTFs are likely acting in cancer cells while others are more likely specific to cells from the tumour microenvironment (most probably immune infiltrating cells). Accordingly, the CpGs whose methylation levels are associated with the expression of the emTFs are predicted to regulate genes enriched for cancer-associated or immune pathways. Finally, we experimentally (i) confirmed the effect of FOXA1 expression on DNA methylation patterns at regions bound by FOXA1 in the MCF-7 breast cancer cell line, and (ii) detected interactions of FOXA1 with TET1 and TET2 proteins both in an *in vitro* setup and at endogenous levels. The *in vitro* GST-pulldown assay of GST fused mTET2 further revealed that the interaction observed between TET2 and FOXA1 is mapped to the C-terminal end of TET2 (amino acid residues 1388-2002). Altogether, the findings outlined in this study provide evidence supporting the importance of specific pioneer TFs in reshaping the chromatin landscape in cancer patients to rewire gene regulatory networks through local DNA demethylation of *cis*-regulatory regions.

The results highlighted in this report complement previous investigations of the interplay between TF binding and DNA methylation. The high-throughput screening approach developed by Suzuki *et al*. [6] exhibited that some developmental TFs induce binding site-directed DNA demethylation. The screening approach requires to select a set of TFs to test and is based on TF overexpression in specific cell lines, while the emQTL approach allows for the large-scale assessment of TFs from cancer patient material. The emQTL methodology has the potential to highlight the physiological and spatio-temporal context of TFs’ expression in cancer samples. Another study leveraged DNA methylation status to identify transcriptional enhancers in cancer samples and predicted their transcriptional targets through correlation of enhancer activity and neighbouring gene expression [23]. These enhancers were enriched for binding profiles of TFs that were predicted as upstream regulators, whose binding might have an effect on DNA methylation [23]. Among these predicted upstream regulators, multiple predictions are in agreement with the emTFs identified here: FOXA1 and GATA3 in BRCA, FOXA2 in UCEC, RUNX1 in KIRP, and SOX2 and TP63 in LUSC cohorts. Moreover, we found that emCpGs were predominantly located in intronic and intergenic regions, which are often associated with enhancers. This observation combined with previous predictions from [23] suggests that the emTFs are likely to drive demethylation at numerous enhancer regions.

We considered in this study a collection of TFBSs with experimental and computational support for direct TF-DNA interactions, which are stored in the UniBind database [25]. This collection was obtained through the uniform processing of thousands of ChIP-seq experiments from diverse cell types / tissues and conditions. We acknowledge that some TFBSs might not be functional in the cancer cells or cell type of origin analyzed here. Nevertheless, they provide the necessary background for large-scale analysis and these regions have been identified as TF-bound in biological contexts. Furthermore, the TFBSs stored in UniBind represent evolutionarily conserved elements [25] and harbour similar mutational load than protein-coding exons (using TCGA somatic mutation data), supporting their functional relevance [55].

In the emQTL analysis performed in this study, TF expression was used as a surrogate to the capacity of TFs to bind their TFBSs. We considered ∼400 bp surrounding TFBSs (± 200 bp) to assess the local effect of TF binding on DNA methylation following [6] where the authors estimated that TF-induced DNA demethylation was local to the TFBSs with a range of a few hundred base pairs (from ∼106 to ∼320 bp). We acknowledge that the RNA expression of a TF might not always relate to its capacity to bind its TFBSs. Nevertheless, increasing TF concentration is related to the capacity of a TF to bind more DNA segments with distinct affinities [56]. Furthermore, we acknowledge that the regulatory activity of TFs goes beyond what can be estimated through their level of transcription. Indeed, several post-translational modifications (PTMs), such as phosphorylation, SUMOylation, ubiquitination, acetylation glycosylation, and methylation are regarded as one type of regulatory mechanism controlling the activity of TFs [57–60]. Unfortunately, capturing PTM information for all TFs in cancer samples is intractable. Past efforts aimed at classifying TFs based on their functional features such as their involvement in signal response versus cell specific developmental function [61]. It is noteworthy that several of the 13 emTF appear to be in the developmental group. For instance, GATA3 is required for the T-helper 2 (Th2) differentiation process (reviewed in [62]); C/EBPB in adipocyte differentiation [63,64]; PBX3 is a homeodomain protein, which are known to be important for human developmental processes [65]; RUNX1 and RUNX3 have a primary role in the development of all hematopoietic cell types [66]; and FLI1 plays an essential role in embryogenesis, vascular development, and megakaryopoiesis [67,68]. As expected, this may indicate that our methodology selects for TFs whose expressions are of key importance for their function.

We further recognize that, as we previously observed for emQTLs in general [26], emTFs are not specific to cancer cells. Indeed, TCGA data were obtained from populations of heterogeneous cancer cells and cells from the tumour microenvironment. Nevertheless, we argue that the heterogeneity of the cells provides the appropriate means to perform correlation analysis such as emQTLs. Furthermore, this strategy provided us with the opportunity to capture signals coming from both cancer cells and immune cells, which could be disentangled through the assessment of tumour purity in TCGA samples.

Several studies previously proposed a model where pioneer TFs remodel the chromatin landscape through increased accessibility followed by DNA methylation loss priming inaccessible enhancers during cell fate transitions (reviewed in [18]). Barnett *et al*. validated this model by profiling DNA methylation and chromatin accessibility at the same time from a single DNA fragment, where they differentiate THP-1 cells into naive M(-) macrophages. They reported that along the enhancer regulation continuum during differentiation of THP-1 cells, loss of DNA methylation is necessary for cell fate determination [18]. Similarly, Reizel *et al*. demonstrated that FOXA1 and FOXA2 TFs are responsible for DNA demethylation at tissue-specific enhancers during liver development, likely through the recruitment of TET2/3 enzymes [69]. Furthermore, pioneer TFs act as developmental factors by controlling key regulatory processes leading to cell identity changes. With the predicted emTFs strongly enriched for pioneer function, we hypothesize that they trigger the aberrant activation of developmental cis-regulatory regions leading to cell identity transitions during carcinogenesis. This hypothesis is in agreement with previous observations of architectural protein- and pioneer TF-mediated chromatin rearrangements that lead to reactivation of embryonic gene expression signatures occuring during cancer (reviewed in [31]).

Several TFs have been reported with a protective role against *de novo* DNA methylation. These TFs include SPI1 [42,43,70–72], YY1, NRF1 [45,71], GABPA, NF-YA [71], CTCF [44], and KLF9 [73]. In line with these reports, we found consistent enrichment for TFBSs associated with these TFs proximal to CpGs harbouring constant hypomethylation across patients, despite the presence of TFBSs for emTFs. On the contrary, regions surrounding emCpGs were enriched for TFBSs associated with the emTFs. This enrichment suggests that several TFBSs for the same emTF colocalize in regulatory regions, which is a signature of homotypic clusters of TFBSs [74]. These homotypic clusters have been described as key components of human promoters and enhancers and have been found to be enriched in developmental enhancers [74]. The expression of the majority of the emTFs exhibited anti-correlation with CpG methylation close to their TFBSs, indicating that these emTFs are likely inducing local DNA demethylation. This is in agreement with previous studies that reported RUNX1 [7,75], RUNX3 [6], SPI1 [6,20,75], BHLHE40 [75], and FOXA1 [22] to induce binding-site directed DNA demethylation. Altogether, these observations provide further evidence for the involvement of emTFs in the specific transcriptional activation of developmental cis-regulatory regions in cancers.

We also captured positively correlated emCpGs, wherein higher TF expression is associated with higher DNA methylation near their TFBS, although in smaller proportions. It is noteworthy that recent work from diverse model systems suggests that 5mC might not always act as a dominant repressive mechanism and that hypermethylated promoters and enhancers can be permissive to transcription *in vivo* and *in vitro* (Reviewed in [76]).

The emQTL analysis did not examine specifically the CpGs lying within TFBSs but rather considered CpGs located at most 200 bp regions away from the TFBSs. As a consequence, the impact of DNA methylation at the TFBSs was not specifically addressed. From the 13 emTFs predicted, eight have either previously been shown *in vitro* to prefer binding methylated sites [77] or recognize binding motifs that do not contain CpGs (GATA3, SOX2, PBX3, CEBPB, FOXA1, FOXA2, SPI1, and TP63). These characteristics provide an advantage for these TFs to act as pioneer factors since their binding would not be precluded by methylation in close chromatin regions. On the contrary, *in vitro* evidence suggests that the five remaining TFs (ETS1, BHLHE40, FLI1, RUNX1, and RUNX3) do not bind, or more weakly, to methylated sites [77]. Nevertheless, the inhibition of binding via DNA methylation detected *in vitro* is not always observed *in vivo* or can be restricted to some genomic regions [78–80]. Supporting evidence for pioneer function has been reported for FLI1 [37,81,82], RUNX1, and RUNX3 [83,84]. How the rest of the remaining factors can engage with closed chromatin would require further investigations.

We associated emCpGs with target genes by relying on (i) the STITCHIT database of regulatory elements to gene links [46] or (ii) genomic distance. It is well known that cis-regulatory elements may regulate distal genes, which are not necessarily the closest ones [85]. By prioritizing regulatory elements to gene links from STITCHIT, we aimed to rely on regulatory associations previously observed in a large collection of cell types. As some links between regulatory elements to genes might be false positives and as some links might be cell type-specific, we exclusively kept the CpG-gene pairs exhibiting anti-correlation (between DNA methylation and expression) to refine the associations in a cancer type-specific way.

The functional enrichment analyses for the genes predicted to be targets of emCpGs confirmed that the emQTL signals were likely derived from either cancer cells or tumor-infiltrating cells. Indeed, bulk tumor samples from TCGA that were analyzed in this study represent a mixture of cancer cells and cells from the tumor microenvironment [47]. The correlation between emTF expression and tumor purity in the samples allows for the discrimination between the two types of signals. However, the heterogeneity of cancer cells, which belong to several clonal populations, provides an additional level of complexity that was not considered in this study. Nevertheless, the identified emTFs are likely to play a major role in shaping the chromatin landscape at cis-regulatory regions controlling the transcription of cancer- or immune-related genes, respectively.

The experimental assessment of the effect of FOXA1 expression on DNA methylation in the MCF-7 breast cancer cell line revealed a limited number of FOXA1-bound regions with significant differential methylation. This is in line with a recent study [86] where CRISPR knockout (KO) of FOXA1 or GATA3 in HCC1954 cells followed by whole genome bisulfite sequencing revealed 84 FOXA1 hypermethylated regions around FOXA1 TFBSs and 30 around GATA3 TFBSs. The limited effect detected could be explained by the fact that DNA methylation is a stable epigenetic mark and the dynamic regulation of methylation and demethylation are rather slow processes. Indeed, the investigation of DNA methylation turnover using experimental and theoretical frameworks revealed that it takes from several days to weeks [12]. Furthermore, the efficiency of transient transfection of the FOXA1 expression plasmid might be lower than the siRNA transfection efficiency, which can contribute to the small effect observed. As our experimental set-up subjected the cells to siRNA-mediated KD for 72 hours and to transient rescue of FOXA1-V5 ectopic expression for 24 hours, some longer term effects have been missed.

In summary, we reported an interplay between TF binding and DNA methylation marks where the binding of pioneer TFs at their TFBSs are likely to trigger local DNA demethylation that could lead to carcinogenesis. These results confirm the central role for pioneer TFs in aberrant DNA demethylation patterns in cancers. While we experimentally assessed the effect of a single

TF in a cell line, the predictions outlined in this report could be followed up through experimental validation to assess their capacity to drive methylation patterns and carcinogenesis.

## Materials and Methods

### TCGA RNA-seq and methylation data

We obtained patient RNA-seq and DNA methylation array (Illumina 450k arrays) data collected by TCGA for 19 cancer types (LAML-US, BRCA-US, PRAD-US, LUAD-US, LUSC-US, COAD-US, LIHC-US, HNSC-US, THCA-US, GBM-US, LGG-US, KIRC-US, KIRP-US, UCEC-US, STAD-US, SKCM-US, PAAD-US, CESC-US, and BLCA-US) from the ICGC data portal [24,87]. The number of samples for which both RNA-seq and DNA methylation array data was available is provided in Table S1.

### Transcription factor binding sites

Direct TF-DNA interaction predictions were retrieved from the UniBind database (version 2018) for 231 human TFs [25]. TFBS coordinates were provided using the GRCh38 assembly of the human genome and were converted to the GRCh19 assembly using the UCSC liftOver tool [88].

### emQTL computation

We performed emQTL analyses by computing Spearman correlations between the levels of methylation at CpGs and TF expression levels in each cohort independently using the same methodology as previously described [26] with the eMap R package (version 1.2) [89]. The emQTL computation was restricted to CpGs at most 200 bp away from UniBind TFBSs. Intersections between CpG coordinates and extended TFBS regions were obtained using the BedTools version 2.26.0 [90]. For each cancer type, we only considered CpGs with an interquartile range of methylation beta values > 0.1 for the computation as in [26].

For each TF in each cancer type, we selected the correlated CpGs with a Bonferroni corrected p-value < 0.01. We only further considered TFs significantly correlated with at least 5,000 CpGs for downstream analyses. To assess the enrichment for correlated CpGs close to the TF’s TFBSs, we performed Mann-Whitney U (MWU) tests with the set of considered CpGs in the corresponding cohort as the universe. TF-cancer type pairs were considered significant with a MWU Bonferroni-corrected p-value < 0.01. An overview of the computational workflow is provided in Figure S1A.

### Upset and Venn diagram plots

All upset and Venn diagram plots were obtained using Intervene (version 0.6.4) [91].

### Pioneer and flanking accessibility-associated TFs

We compiled a list of pioneer TFs from the literature [30–39] (Table S4). The list of flanking accessibility-associated TFs were retrieved from [92], where they have been described to be associated with increased flanking accessibility around their motif center in cancer samples from ATAC-seq data. We considered in our study the 29 flanking accessibility-associated TFs that were tested for emQTL in this report. We assessed the significance of the intersection between the list of pioneer TFs (or the list of flanking accessibility-associated TFs) and the emTFs by performing Fisher tests with the Bioconductor GeneOverlap package (version 1.18.0) [93].

### Comparison between emCpGs and non-correlated CpGs

For each emTF-cancer type pair, we assessed the enrichment for TFBSs around the corresponding emCpGs and non-correlated CpGs. We computed differential enrichment of TFBSs between regions of ± 200 bp centered around the emCpGs versus the non-correlated CpGs, and vice-versa. Genomic regions were lifted, using the liftOver tool from UCSC [88], from the GRCh19 genome assembly over to the GRCh38 version, which is the assembly used in UniBind. Differential enrichment of TFBS sets was performed using the *twoSets* subcommand of the UniBind enrichment tool (https://unibind.uio.no/enrichment/; https://bitbucket.org/CBGR/unibind_enrichment/) using the collection of TFBS sets from UniBind version 2018 [25,28]. Specifically, the foreground set of regions corresponded to the regions centered around emCpGs or non-correlated CpGs and the combined set of such regions was used as background.

The %GC distributions at genomic regions centered around emCpGs and non-correlated CpGs were computed by the BedTools *nuc* function.

### Association between emCpGs and target genes

We downloaded the associations between regulatory elements and target genes predicted by STITCHIT from the ENCODE, Roadmap, and Blueprint datasets at https://zenodo.org/record/2547384#.XIK0x-RYZ14. The coordinates of the emCpGs considered for each emTF in each cohort were lifted from GRCh19 over to GRCh38 coordinates and intersected with STITCHIT regulatory elements using the *intersect* subcommand of the BedTools. CpGs lying within the regulatory elements were associated with the corresponding target genes. The CpGs not overlapping with STITCHIT regulatory elements were linked to genes with the nearest TSS using the HOMER *annotatePeaks*.*pl* script [94].

### Genomic distribution of CpGs

We used the *annotatePeaks*.*pl* script from HOMER [94] to compute and plot the genomic distribution of all CpGs investigated (n=376,997) and of the emCpGs in each cancer type (Figure S4).

### Correlation between ATAC signal and methylation at emCpGs

We downloaded the TCGA ATAC-seq data described in [40] from https://gdc.cancer.gov/about-data/publications/ATACseq-AWG. We considered the cancer cohorts with at least 20 samples for which DNA methylation was available in our study. We selected the emCpGs predicted in each cancer type and their surrounding ±200 bp regions and intersected them with the pancancer ATAC-seq peaks provided in [40] using the BedTools *intersect* subcommand. Finally, spearman correlation between the level of methylation at emCpGs and the level of ATAC-seq normalized counts at the underlying peaks were computed in each cancer type.

### Functional enrichment analysis

Genes associated to emTFs in cancer cohorts were submitted to the *clusterProfiler* R package (version 3.12.0) [95] to compute enrichment for gene ontology (GO) biological processes and MSigDB Hallmark sets (the *gmt* file corresponding to the Hallmark set was retrieve from MSigDB v7.4). Redundant enriched GO terms were reduced using the *GOSemSim* R package (version 2.10.0) [96]. For MsigDB Hallmark set enriched terms, we considered the top 10 enriched terms (ranked by Benjamini and Hochberg adjusted p-values < 0.05) per emTF-cancer pairs for drawing the figures. In Figure S16, we considered the GO terms with Benjamini and Hochberg adjusted p-values < 0.05 and plotted the top 5 most enriched terms per emTF-cancer pair. Enrichment plots were produced using the *geom_tile* function from the *ggplot2* R package (version 3.3.3).

### Tumour purity

We downloaded cumulative tumor purity estimates from [47]. For the STAD-US cohort, the cumulative tumor purity was not computed in [47]; we retrieved tumor purity scores for STAD-US samples from the ICGC data portal (dcc.icgc.org/releases/PCAWG/consensus_cnv). Pearson correlations between tumour purity and TF RNA expression were computed using the *stat_cor* R function with the parameter *method*=*”pearson”* using the *ggscatter* function in *ggplot2*.

### Bioinformatics analysis of mTET2 and hTET2 proteins

To assess the interaction between TET2 and FOXA1, we used mTET2 GST fusion protein. To assess the relevance of using mTET2 in the GST pull down assay, we assessed the amino acid sequence conservation between mTET2 isoform 2 (RefSeq ID: NP_001035490) and the human TET2 (hTET2; RefSeq ID: NP_001120680). We visualized the pairwise sequence alignment of the two proteins using the MUSCLE algorithm accessed through Jalview (version 2.11.1.4) [97]. It revealed that mTET2 aligns well with the C-terminal half of hTET2 (pairwise sequence identity = 59.72 %; Figure S20A). To further highlight the conservation of the TET2 proteins between mouse and human at the structural level, we obtained the Protein Data Bank structures corresponding to hTET2 (PDB ID: 4nm6A) and the modeled structure of mTET2 (PDB ID: Q6NO21). We compared the two structures with the *ce align* algorithm implemented in pyMOL version 2.4.2, which is represented in Figure S20B.

### Plasmid construction

The human FOXA1 sequence with RefSeq accession ID NM_004496 was synthesized with a C-terminal V5-tag sequence and obtained in pCIneo vector with NheI and XhoI cloning sites from GenScript. The sequence with the C-terminal V5-tag was transferred to the pEF1neo mammalian expression vector using NheI and SalI. pEF1neo is a vector generated from pCIneo by replacing the CMV promoter with the human EF1-alpha promoter. It generates a mammalian expression vector for FOXA1 as pEF1neo-FOXA1-V5. GST-TET1 fusion protein was made by transferring the full length TET1 sequence into the pGEX-KG vector, which was derived from pGEX-2T as described in [98]. The pGEX-KG vector was first cut with XmaI and was filled in with Klenow (Roche Applied Science) to form blunt end, this was followed by XbaI digestion. N-terminally FLAG- and HA-tagged TET1 from the mammalian expression plasmid pEF1-FH-TET1 (ABCAM) was digested with BmgBI (blunt end cutter) and XbaI. The 6436 bp fragment was then inserted into the XmaI and XbaI digested pGEX-KG vector. *Mus musculus* TET2 (mTET2) clone IMAGE ID 4977050 was obtained from Source Bioscience and PCR cloned using oligos (Tet2fwd: 5’-GGGGACAAGTTTGTACAAAAAAGCAGGCTTAatgccaaatggcagtacagt-3’ and Tet2rev: 5’-GGGGACCACTTTGTACAAGAAAGCTGGGTTtcatacaaatgtgttgtaag-3’) into pDonor221 (Invitrogen Gateway, ThermoFisher) and sequenced. mTET2 was then cloned into pGEX-AB-GAW by LR reaction for recombinant protein expression of GST fused mTET2.

### Cell cultures and siRNA and plasmid transfections

MCF-7 cells (ATCC® HTB-22™ *Homo sapiens*, epithelial, mammary gland, breast; derived from metastatic site: pleural effusion, adenocarcinoma) were maintained in RPMI-1640-GlutaMAX supplement medium supplemented with 10% FCS (fetal calf serum) and 1% PS (penicillin/streptomycin), and were grown at 37 °C and 5% CO_2_.

We performed siRNA mediated KD of endogenous FOXA1 from MCF-7 cells stably expressing either pEF1neo-V5 or pEF1neo-FOXA1-V5 using custom synthesized siFOXA1 from Qiagen. The siRNA sequences that target the 3’-UTR of FOXA1 were described in [99]. For control transfections, we used the AllStars Negative Control siRNA (cat.no 1027281, Qiagen). Both siCtrl and siFOXA1 at a concentration of 10 µM were delivered to cells using the lullaby siRNA transfection reagent (OZ biosciences). Specifically, cells were seeded at a density of 10^5^ cells in 6 well plates 24 hours prior to siRNA transfection. The next day, the media was changed and siRNAs were delivered using lullaby siRNA transfection reagent. The cells were subjected to siRNA mediated KD for 72 hours at 37 °C and 5% CO_2_ before they were harvested for DNA isolation. Transient rescue of the endogenous KD was made 2 days post siRNA transfection by delivering 2.5 µg of pEF1neo-FOXA1-V5 plasmid with the lullaby transfection reagent for 24 hours.

For GST pull-down assay, COS-1 cells were transiently transfected with either 5 or 10 μg of each of pCIneo-FOXA1-V5 and pCIneo-V5 plasmids using lipofectamine 3000 Reagent (Invitrogen).

### Methylation array profiling in MCF-7 cells and bioinformatics analysis

Genomic DNA from MCF-7 cells transfected with either siCtrl or siFOXA1 and MCF-7 cells subjected to siFOXA1 mediated endogenous KD and rescue with exogenous FOXA1-V5 in three biological replicates was isolated using NucleoSpin® Tissue genomic DNA isolation kit (Machery-Nagel). From each sample, 45 µl genomic DNA amounting to 500 ng concentration was delivered to the Genomics core facility at Oslo University hospital, where EPIC array profiling was performed. Bisulfite-converted DNA was amplified, fragmented, and hybridised to Illumina Infinium Human Methylation 850K Beadchip using standard Illumina protocol.

EPIC array methylation data in IDAT format were normalized with the *minfi* (version 1.36.0) R package [100] using the within array Noob function followed by quantile normalization as recommended by shinyÉpico [101]. M-values were obtained from the normalized -values using *minfi*. Contrasts of M-values were computed using the *limma* (version 3.46.0) R package between control and KD replicates with the *limma::arrayWeights* option to mitigate the influence of the arrays. The computed raw p-values from the *limma* fit were provided to the *mCSEATest* function of the *mCSEA* R package (version 1.10.0) to compute differentially methylated regions considering FOXA1 ChIP-seq peaks. FOXA1 ChIP-seq peaks were retrieved from the ReMap 2020 database [102] considering ChIP-seq experiments performed in MCF-7 cells without target or biotype modification.

To draw the heatmap provided in Figure 4A, we considered all CpGs in the EPIC array lying within the identified DMRs (n=431). For each CpG, we computed the average of the -values across the three control replicates. The average value was subtracted from the -value of each CpG in each of the nine samples. Finally, we filtered out the resulting values *vals* that satisfied - 0.05 < *vals* < 0.05. The remaining values associated with 228 CpGs were plotted in Figure 4A using the *pheatmap* R package (version 1.0.12).

### MCF-7 nuclear extract preparation

Nuclear extracts from MCF-7 cells were prepared as described in [103] with a slight modification. To disrupt the cytoplasmic membrane, in addition to douncing, detergent was used by supplementing buffer A with 0.05% NP-40.

### Antibodies

For western blot (WB) validation of positive clones and endogenous KDs, we used the following primary antibodies: mouse anti-V5 monoclonal antibody (46-0705, Invitrogen), rabbit anti-FOXA1 M2 polyclonal antibody (GTX100308, Gene-Tex), and mouse anti GAPDH monoclonal antibody (AM4300, Invitrogen). We used the following secondary antibodies for WB: anti-mouse IRDye® 680 RD (925-68072, LICOR) and anti-mouse IRDye 800 CW (925-32213, LI COR).

For endogenous immunoprecipitation, we used the following antibodies: anti-FOXA1 M2 rabbit polyclonal antibody (GTX100308, Gene-Tex), anti-TET1 mouse monoclonal antibody (GTX627420, Gene-Tex), anti-TET2 Rabbit monoclonal antibody (D6B9Y, cell signalling technologies), normal rabbit IgG (2729S, cell signalling technologies), normal mouse IgG (sc-2025, Santa Cruz), and protein G Dynabeads (10004D, Invitrogen).

### GST-pulldown and immunoprecipitation assays

The GST fusion proteins and GST were expressed and isolated as described in [104]. Total cell lysates from COS-1 cells 24 hr post transfection were prepared using 300 μl of KAc interaction buffer (Roche Applied Science). GST fusion proteins were bound to glutathione-Sepharose beads (GE Healthcare) by rotating in binding buffer (50 mM Tris HCl pH 8.0, 150 mM NaCl, 5 mM EDTA, 1% Triton X-100, 1 mM Dithiothreitol (DTT) and 1× Complete protease inhibitor cocktail at 4 °C for 1 hr prior to pull-down. The pre-bound fusion proteins were then incubated for 1 hr at 4 °C with whole cell lysate obtained from transfected COS-1 cells. The beads were washed 3× in 500 μl of KAc interaction buffer. The bound proteins were eluted in 40 μl of 3× SDS loading buffer at 95 °C for 10 min and detected using immunoblotting after SDS-PAGEPAGE separation on a 4-15% SDS-PAA gel and western blotting.

Immunoprecipitation at endogenous level of FOXA1, TET1, and TET2 was obtained by incubating rabbit anti-FOXA1 polyclonal, mouse anti-TET1 monoclonal, and rabbit anti-TET2 monoclonal antibodies, respectively coupled with protein G Dynabeads (Invitrogen) with nuclear extract derived from MCF-7 cells for 2 hours, with rotation at 4 °C. As negative controls, mouse or rabbit normal IgG coupled with protein G Dynabeads were used. Prior to incubation, we washed the beads once with 1× PBS supplemented with 0.03 µg BSA and further blocked them with 0.03 µg BSA in 1× PBS for 10 minutes with rotation. We then washed the beads twice with 400 µl lysis buffer (20mM HEPES, 10% Glycerol, 0,05%NP-40, 1,5 mM MgCl2, 150 mM KAc, and 1 mM DTT supplemented with 5× Complete protease inhibitor cocktail). Each wash was performed for 5 minutes with rotation at 4 °C. The bound proteins were eluted with a 20 μl 3× SDS loading buffer at 95 °C for 10 min. After SDS-PAGE separation on a 4-15% SDS-PAA gel, the proteins were detected with western blot using a OdysseyCLX (LI COR).

## Data and code availability

The code used to generate the emQTL analysis and the processing of the methylation array data is available at https://bitbucket.org/CBGR/tfme/src/master/. The raw and processed methylation data are available on GEO with the accession number GSE174008.

## Acknowledgements

The methylation array was performed at the Genomics Core Facility, Oslo University Hospital (http://oslo.genomics.no/). We thank Prof. Reidun Aalen for providing the pGEX-AB-GAW vector. We thank Ieva Rauluseviciute for thoroughly reading the manuscript and providing valuable comments, Rafael Riudavets Puig and Vipin Kumar for reading the manuscript and suggestions on the analysis, the Mathelier and Kuijjer groups for insightful discussions, Johannes Landskron and his team for help with mycoplasma test of the cells, Sebastian Waszak for his help with the PCAWG tumor purity data, Georgios Magklaras, Harold Gutch, and the NCMM IT team for IT support, and Ingrid Kjelsvik for administrative support.

## Funding

Research Council of Norway [187615], Helse Sør-Øst, and University of Oslo through the Centre for Molecular Medicine Norway (NCMM) (to Mathelier group); Research Council of Norway [288404 Mathelier group]; Norwegian Cancer Society [197884 to RBL and Mathelier group]. Research Council of Norway through its Centres of Excellence funding scheme [262652 to RE].

## Competing interests

None declared.

## Notes

### Competing Interest Statement

The authors have declared no competing interest.

### Summary of Updates

In this revision, the main modifications are new analyses of ATAC-seq data, new results from GST pull down experiments, and new paragraphs in the Discussion section to provide an even more extensive context to our work.

## References

1. Jaenisch R, Bird A. Epigenetic regulation of gene expression: how the genome integrates intrinsic and environmental signals. Nat Genet. 2003;33 Suppl:245–54.

2. Kouzarides T. Chromatin modifications and their function. Cell. 2007;128:693–705.

3. Sood AJ, Viner C, Hoffman MM. DNAmod: the DNA modification database. J Cheminform. 2019;11:30.

4. Breiling A, Lyko F. Epigenetic regulatory functions of DNA modifications: 5-methylcytosine and beyond. Epigenetics Chromatin. 2015;8:24.

5. Curradi M, Izzo A, Badaracco G, Landsberger N. Molecular mechanisms of gene silencing mediated by DNA methylation. Mol Cell Biol. 2002;22:3157–73.

6. Suzuki T, Maeda S, Furuhata E, Shimizu Y, Nishimura H, Kishima M, et al. A screening system to identify transcription factors that induce binding site-directed DNA demethylation. Epigenetics Chromatin. 2017;10:60.

7. Suzuki T, Shimizu Y, Furuhata E, Maeda S, Kishima M, Nishimura H, et al. RUNX1 regulates site specificity of DNA demethylation by recruitment of DNA demethylation machineries in hematopoietic cells. Blood Adv. 2017;1:1699–711.

8. Zhu H, Wang G, Qian J. Transcription factors as readers and effectors of DNA methylation. Nat Rev Genet. 2016;17:551–65.

9. Rasmussen KD, Helin K. Role of TET enzymes in DNA methylation, development, and cancer. Genes Dev. 2016;30:733–50.

10. Williams K, Christensen J, Pedersen MT, Johansen JV, Cloos PAC, Rappsilber J, et al. TET1 and hydroxymethylcytosine in transcription and DNA methylation fidelity. Nature. 2011;473:343–8.

11. Laisné M, Gupta N, Kirsh O, Pradhan S, Defossez P-A. Mechanisms of DNA Methyltransferase Recruitment in Mammals. Genes. 2018;9.

12. Ginno PA, Gaidatzis D, Feldmann A, Hoerner L, Imanci D, Burger L, et al. A genome-scale map of DNA methylation turnover identifies site-specific dependencies of DNMT and TET activity. Nat Commun. 2020;11:2680.

13. Lambert SA, Jolma A, Campitelli LF, Das PK, Yin Y, Albu M, et al. The Human Transcription Factors. Cell. 2018;172:650–65.

14. Reiter F, Wienerroither S, Stark A. Combinatorial function of transcription factors and cofactors. Curr Opin Genet Dev. 2017;43:73–81.

15. Zaret KS, Carroll JS. Pioneer transcription factors: establishing competence for gene expression. Genes Dev. 2011;25:2227–41.

16. Young RA. Control of the embryonic stem cell state. Cell. 2011;144:940–54.

17. Jozwik KM, Carroll JS. Pioneer factors in hormone-dependent cancers. Nat Rev Cancer. 2012;12:381–5.

18. Barnett KR, Decato BE, Scott TJ, Hansen TJ, Chen B, Attalla J, et al. ATAC-Me Captures Prolonged DNA Methylation of Dynamic Chromatin Accessibility Loci during Cell Fate Transitions. Mol Cell. 2020.

19. Di Croce L, Raker VA, Corsaro M, Fazi F, Fanelli M, Faretta M, et al. Methyltransferase recruitment and DNA hypermethylation of target promoters by an oncogenic transcription factor. Science. 2002;295:1079–82.

20. de la Rica L, Rodríguez-Ubreva J, García M, Islam ABMMK, Urquiza JM, Hernando H, et al. PU.1 target genes undergo Tet2-coupled demethylation and DNMT3b-mediated methylation in monocyte-to-osteoclast differentiation. Genome Biol. 2013;14:R99.

21. Yang YA, Zhao JC, Fong K-W, Kim J, Li S, Song C, et al. FOXA1 potentiates lineage-specific enhancer activation through modulating TET1 expression and function. Nucleic Acids Res. 2016;44:8153–64.

22. Vanzan L, Soldati H, Ythier V, Anand S, Braun SMG, Francis N, et al. High throughput screening identifies SOX2 as a super pioneer factor that inhibits DNA methylation maintenance at its binding sites. Nat Commun. 2021;12:3337.

23. Yao L, Shen H, Laird PW, Farnham PJ, Berman BP. Inferring regulatory element landscapes and transcription factor networks from cancer methylomes. Genome Biol. 2015;16:105.

24. Cancer Genome Atlas Research Network, Weinstein JN, Collisson EA, Mills GB, Shaw KRM, Ozenberger BA, et al. The Cancer Genome Atlas Pan-Cancer analysis project. Nat Genet. 2013;45:1113–20.

25. Gheorghe M, Sandve GK, Khan A, Chèneby J, Ballester B, Mathelier A. A map of direct TF-DNA interactions in the human genome. Nucleic Acids Res. 2019; 4:e21.

26. Fleischer T, Tekpli X, Mathelier A, Wang S, Nebdal D, Dhakal HP, et al. DNA methylation at enhancers identifies distinct breast cancer lineages. Nat Commun. 2017;8:1379.

27. Sandoval J, Heyn H, Moran S, Serra-Musach J, Pujana MA, Bibikova M, et al. Validation of a DNA methylation microarray for 450,000 CpG sites in the human genome. Epigenetics. 2011;6:692–702.

28. Puig RR, Boddie P, Khan A, Castro-Mondragon JA, Mathelier A. UniBind: maps of high-confidence direct TF-DNA interactions across nine species. BMC Genomics. 2021;22:482.

29. Jones PA, Baylin SB. The epigenomics of cancer. Cell. 2007;128:683–92.

30. Iwafuchi-Doi M, Zaret KS. Pioneer transcription factors in cell reprogramming. Genes Dev. 2014;28:2679–92.

31. Dobersch S, Rubio K, Barreto G. Pioneer Factors and Architectural Proteins Mediating Embryonic Expression Signatures in Cancer. Trends Mol Med. 2019;25:287–302.

32. Vernimmen D, Bickmore WA. The Hierarchy of Transcriptional Activation: From Enhancer to Promoter. Trends Genet. 2015;31:696–708.

33. Fuglerud BM, Lemma RB, Wanichawan P, Sundaram AYM, Eskeland R, Gabrielsen OS. A c-Myb mutant causes deregulated differentiation due to impaired histone binding and abrogated pioneer factor function. Nucleic Acids Res. 2017;45:7681–96.

34. Hoogenkamp M, Lichtinger M, Krysinska H, Lancrin C, Clarke D, Williamson A, et al. Early chromatin unfolding by RUNX1: a molecular explanation for differential requirements during specification versus maintenance of the hematopoietic gene expression program. Blood. 2009;114:299–309.

35. Wang D, Diao H, Getzler AJ, Rogal W, Frederick MA, Milner J, et al. The Transcription Factor Runx3 Establishes Chromatin Accessibility of cis-Regulatory Landscapes that Drive Memory Cytotoxic T Lymphocyte Formation. Immunity. 04 17, 2018;48:659–74.e6.

36. Lee J-W, Kim D-M, Jang J-W, Park T-G, Song S-H, Lee Y-S, et al. RUNX3 regulates cell cycle-dependent chromatin dynamics by functioning as a pioneer factor of the restriction-point. Nat Commun. 2019;10:1897.

37. Riggi N, Knoechel B, Gillespie SM, Rheinbay E, Boulay G, Suvà ML, et al. EWS-FLI1 utilizes divergent chromatin remodeling mechanisms to directly activate or repress enhancer elements in Ewing sarcoma. Cancer Cell. 2014;26:668–81.

38. Pang Z-Y, Wei Y-T, Shang M-Y, Li S, Li Y, Jin Q-X, et al. Leptin-elicited PBX3 confers letrozole resistance in breast cancer. Endocr Relat Cancer. 2021;28(3), 173–189.

39. Plachetka A, Chayka O, Wilczek C, Melnik S, Bonifer C, Klempnauer K-H. C/EBPbeta induces chromatin opening at a cell-type-specific enhancer. Mol Cell Biol. 2008;28:2102–12.

40. Corces MR, Granja JM, Shams S, Louie BH, Seoane JA, Zhou W, et al. The chromatin accessibility landscape of primary human cancers. Science. 2018;362.

41. Baek S, Goldstein I, Hager GL. Bivariate Genomic Footprinting Detects Changes in Transcription Factor Activity. Cell Rep. 2017;19:1710–22.

42. Brandeis M, Frank D, Keshet I, Siegfried Z, Mendelsohn M, Names A, et al. Spl elements protect a CpG island from de novo methylation. Nature. 1994;371:435–8.

43. Macleod D, Charlton J, Mullins J, Bird AP. Sp1 sites in the mouse aprt gene promoter are required to prevent methylation of the CpG island. Genes Dev. 1994;8:2282–92.

44. Pant V, Mariano P, Kanduri C, Mattsson A. The nucleotides responsible for the direct physical contact between the chromatin insulator protein CTCF and the H19 imprinting control region manifest parent of origin-specific long-distance insulation and methylation-free domains. Genes Dev. 2003 Mar 1;17(5):586–90.

45. Gebhard C, Benner C, Ehrich M, Schwarzfischer L, Schilling E, Klug M, et al. General transcription factor binding at CpG islands in normal cells correlates with resistance to de novo DNA methylation in cancer cells. Cancer Res. 2010;70:1398–407.

46. Schmidt F, Marx A, Hebel M, Wegner M, Baumgarten N. Integrative analysis of epigenetics data identifies gene-specific regulatory elements. bioRxiv. biorxiv.org; 2019; doi.org/10.1101/585125

47. Aran D, Sirota M, Butte AJ. Systematic pan-cancer analysis of tumour purity. Nat Commun. 2015;6:8971.

48. Liberzon A, Birger C, Thorvaldsdóttir H, Ghandi M, Mesirov JP, Tamayo P. The Molecular Signatures Database (MSigDB) hallmark gene set collection. Cell Syst. 2015;1:417–25.

49. Iwafuchi-Doi M, Zaret KS. Cell fate control by pioneer transcription factors. Development. 2016;143:1833–7.

50. Morris SA. Direct lineage reprogramming via pioneer factors; a detour through developmental gene regulatory networks. Development. 2016;143:2696–705.

51. Martorell-Marugán J, González-Rumayor V, Carmona-Sáez P. mCSEA: detecting subtle differentially methylated regions. Bioinformatics. 2019;35:3257–62.

52. Rae JM, Johnson MD, Scheys JO, Cordero KE, Larios JM, Lippman ME. GREB 1 is a critical regulator of hormone dependent breast cancer growth. Breast Cancer Res Treat. Springer Science and Business Media LLC; 2005;92:141–9.

53. Prest SJ, May FEB, Westley BR. The estrogen-regulated protein, TFF1, stimulates migration of human breast cancer cells. FASEB J. Wiley; 2002;16:592–4.

54. Cantor S, Drapkin R, Zhang F, Lin Y, Han J, Pamidi S, et al. The BRCA1-associated protein BACH1 is a DNA helicase targeted by clinically relevant inactivating mutations. Proc Natl Acad Sci U S A. 2004;101:2357–62.

55. Castro-Mondragon JA, Aure MR, Lingærde OC. Cis-regulatory mutations associate with transcriptional and post-transcriptional deregulation of the gene regulatory program in cancers. bioRxiv 2020; doi.org/10.1101/2020.06.25.170738

56. Kribelbauer JF, Rastogi C, Bussemaker HJ, Mann RS. Low-Affinity Binding Sites and the Transcription Factor Specificity Paradox in Eukaryotes. Annu Rev Cell Dev Biol. 2019;35:357–79.

57. Walsh CT, Garneau-Tsodikova S, Gatto GJ Jr. Protein posttranslational modifications: the chemistry of proteome diversifications. Angew Chem Int Ed Engl. 2005;44:7342–72.

58. Geiss-Friedlander R, Melchior F. Concepts in sumoylation: a decade on. Nat Rev Mol Cell Biol. 2007;8:947–56.

59. Benayoun BA, Veitia RA. A post-translational modification code for transcription factors: sorting through a sea of signals. Trends Cell Biol. 2009;19:189–97.

60. Filtz TM, Vogel WK, Leid M. Regulation of transcription factor activity by interconnected post-translational modifications. Trends Pharmacol Sci. 2014;35:76–85.

61. Brivanlou AH, Darnell JE Jr. Signal transduction and the control of gene expression. Science. 2002;295:813–8.

62. Zhu J. Transcriptional regulation of Th2 cell differentiation. Immunology & Cell Biology. 2010. p. 244–9.

63. Hishida T, Nishizuka M, Osada S, Imagawa M. The role of C/EBPdelta in the early stages of adipogenesis. Biochimie. 2009;91:654–7.

64. Siersbæk R, Mandrup S. Transcriptional networks controlling adipocyte differentiation. Cold Spring Harb Symp Quant Biol. 2011;76:247–55.

65. Bürglin TR. Homeobox Genes. In: Maloy S, Hughes K, editors. Brenner’s Encyclopedia of Genetics (Second Edition). San Diego: Academic Press; 2013. p. 503–8.

66. de Bruijn M, Dzierzak E. Runx transcription factors in the development and function of the definitive hematopoietic system. Blood. 2017;129:2061–9.

67. Wang C, Sample KM, Gajendran B, Kapranov P, Liu W, Hu A, et al. FLI1 Induces Megakaryopoiesis Gene Expression Through WAS/WIP-Dependent and Independent Mechanisms; Implications for Wiskott-Aldrich Syndrome. Front Immunol. 2021;12:607836.

68. Hart A, Melet F, Grossfeld P, Chien K, Jones C, Tunnacliffe A, et al. Fli-1 is required for murine vascular and megakaryocytic development and is hemizygously deleted in patients with thrombocytopenia. Immunity. 2000;13:167–77.

69. Reizel Y, Morgan A, Gao L, Schug J, Mukherjee S, García MF, et al. FoxA-dependent demethylation of DNA initiates epigenetic memory of cellular identity. Dev Cell. 2021;56:602–12.e4.

70. Boumber YA, Kondo Y, Chen X, Shen L, Guo Y, Tellez C, et al. An Sp1/Sp3 Binding Polymorphism Confers Methylation Protection. PLoS Genet. 2008;4:e1000162.

71. Berman BP, Weisenberger DJ, Aman JF, Hinoue T, Ramjan Z, Liu Y, et al. Regions of focal DNA hypermethylation and long-range hypomethylation in colorectal cancer coincide with nuclear lamina--associated domains. Nat Genet. Nature Publishing Group; 2012;44:40.

72. Han L, Lin IG, Hsieh CL. Protein binding protects sites on stable episomes and in the chromosome from de novo methylation. Mol Cell Biol. 2001;21:3416–24.

73. Li J, Abe K, Milanesi A, Liu Y-Y, Brent GA. Thyroid Hormone Protects Primary Cortical Neurons Exposed to Hypoxia by Reducing DNA Methylation and Apoptosis. Endocrinology. 2019;160:2243–56.

74. Gotea V, Visel A, Westlund JM, Nobrega MA, Pennacchio LA, Ovcharenko I. Homotypic clusters of transcription factor binding sites are a key component of human promoters and enhancers. Genome Res. 2010;20:565–77.

75. Martin-Trujillo A, Patel N, Richter F, Jadhav B, Garg P, Morton SU, et al. Rare genetic variation at transcription factor binding sites modulates local DNA methylation profiles. PLoS Genet. 2020;16:e1009189.

76. Angeloni A, Bogdanovic O. Enhancer DNA methylation: implications for gene regulation. Essays Biochem. 2019;63:707–15.

77. Yin Y, Morgunova E, Jolma A, Kaasinen E, Sahu B, Khund-Sayeed S, et al. Impact of cytosine methylation on DNA binding specificities of human transcription factors. Science. 2017;356.

78. Domcke S, Bardet AF, Adrian Ginno P, Hartl D, Burger L, Schübeler D. Competition between DNA methylation and transcription factors determines binding of NRF1. Nature. 2015;528:575–9.

79. Becker PB, Ruppert S, Schütz G. Genomic footprinting reveals cell type-specific DNA binding of ubiquitous factors. Cell. 1987;51:435–43.

80. Stadler MB, Murr R, Burger L, Ivanek R, Lienert F, Schöler A, et al. DNA-binding factors shape the mouse methylome at distal regulatory regions. Nature. 2011. p. 490–5.

81. Martens JHA, Mandoli A, Simmer F, Wierenga B-J, Saeed S, Singh AA, et al. ERG and FLI1 binding sites demarcate targets for aberrant epigenetic regulation by AML1-ETO in acute myeloid leukemia. Blood. 2012;120:4038–48.

82. Giraud G, Kolovos P, Boltsis I, van Staalduinen J, Guyot B, Weiss-Gayet M, et al. Interplay between FLI-1 and the LDB1 complex in murine erythroleukemia cells and during megakaryopoiesis. iScience. 2021;24:102210.

83. Mevel R, Draper JE, Lie-A-Ling M, Kouskoff V, Lacaud G. RUNX transcription factors: orchestrators of development. Development. 2019;146.

84. Hass MR, Brissette D, Parameswaran S, Pujato M, Donmez O, Kottyan LC, et al. Runx1 shapes the chromatin landscape via a cascade of direct and indirect targets. PLoS Genet. 2021;17:e1009574.

85. Schoenfelder S, Fraser P. Long-range enhancer–promoter contacts in gene expression control. Nat Rev Genet. Nature Publishing Group; 2019;20:437–55.

86. Detilleux D, Spill YG, Balaramane D, Weber M, Bardet AF. Focal DNA hypo-methylation in cancer is mediated by transcription factors binding. bioRxiv. 2021; doi.org/10.1101/2021.04.20.440687

87. Zhang J, Bajari R, Andric D, Gerthoffert F, Lepsa A, Nahal-Bose H, et al. The International Cancer Genome Consortium Data Portal. Nat Biotechnol. 2019;37:367–9.

88. Hinrichs AS, Karolchik D, Baertsch R, Barber GP, Bejerano G, Clawson H, et al. The UCSC Genome Browser Database: update 2006. Nucleic Acids Res. 2006;34:D590–8.

89. Sun W. eMap: map gene expression qtl. R package version 1.2. 2010.

90. Quinlan AR, Hall IM. BEDTools: a flexible suite of utilities for comparing genomic features. Bioinformatics. 2010;26:841–2.

91. Khan A, Mathelier A. Intervene: a tool for intersection and visualization of multiple gene or genomic region sets. BMC Bioinformatics. 2017;18:287.

92. The chromatin accessibility landscape of primary human cancers. Science 2019; http://science.sciencemag.org/content/362/6413/eaav1898

93. Li Shen, Mount Sinai. GeneOverlap. Bioconductor; 2017.

94. Heinz S, Benner C, Spann N, Bertolino E, Lin YC, Laslo P, et al. Simple combinations of lineage-determining transcription factors prime cis-regulatory elements required for macrophage and B cell identities. Mol Cell. 2010;38:576–89.

95. Yu G, Wang L-G, Han Y, He Q-Y. clusterProfiler: an R package for comparing biological themes among gene clusters. OMICS. 2012;16:284–7.

96. Yu G, Li F, Qin Y, Bo X, Wu Y, Wang S. GOSemSim: an R package for measuring semantic similarity among GO terms and gene products. Bioinformatics. 2010. p. 976–8.

97. Waterhouse AM, Procter JB, Martin DMA, Clamp M, Barton GJ. Jalview Version 2--a multiple sequence alignment editor and analysis workbench. Bioinformatics. 2009;25:1189–91.

98. Guan KL, Dixon JE. Eukaryotic proteins expressed in Escherichia coli: an improved thrombin cleavage and purification procedure of fusion proteins with glutathione S-transferase. Anal Biochem. 1991;192:262–7.

99. Carroll JS, Liu XS, Brodsky AS, Li W, Meyer CA, Szary AJ, et al. Chromosome-wide mapping of estrogen receptor binding reveals long-range regulation requiring the forkhead protein FoxA1. Cell. 2005;122:33–43.

100. Aryee MJ, Jaffe AE, Corrada-Bravo H, Ladd-Acosta C, Feinberg AP, Hansen KD, et al. Minfi: a flexible and comprehensive Bioconductor package for the analysis of Infinium DNA methylation microarrays. Bioinformatics. 2014;30:1363–9.

101. Morante-Palacios O, Ballestar E. shinyÉPICo: A graphical pipeline to analyze Illumina DNA methylation arrays. Bioinformatics. 2021; http://dx.doi.org/10.1093/bioinformatics/btaa1095

102. Chèneby J, Ménétrier Z, Mestdagh M, Rosnet T, Douida A, Rhalloussi W, et al. ReMap 2020: a database of regulatory regions from an integrative analysis of Human and Arabidopsis DNA-binding sequencing experiments. Nucleic Acids Res. 2020;48:D180–8.

103. Rodríguez-Castañeda F, Lemma RB, Cuervo I, Bengtsen M, Moen LM, Ledsaak M, et al. The SUMO protease SENP1 and the chromatin remodeler CHD3 interact and jointly affect chromatin accessibility and gene expression. J Biol Chem. 2018;293:15439–54.

104. Dahle È, Bakke O, Gabrielsen OS. c-Myb associates with PML in nuclear bodies in hematopoietic cells. Exp Cell Res. 2004;297:118–26.

