## Supplementary Information for "Pioneer transcription factors are associated with the modulation of DNA methylation patterns across cancers"

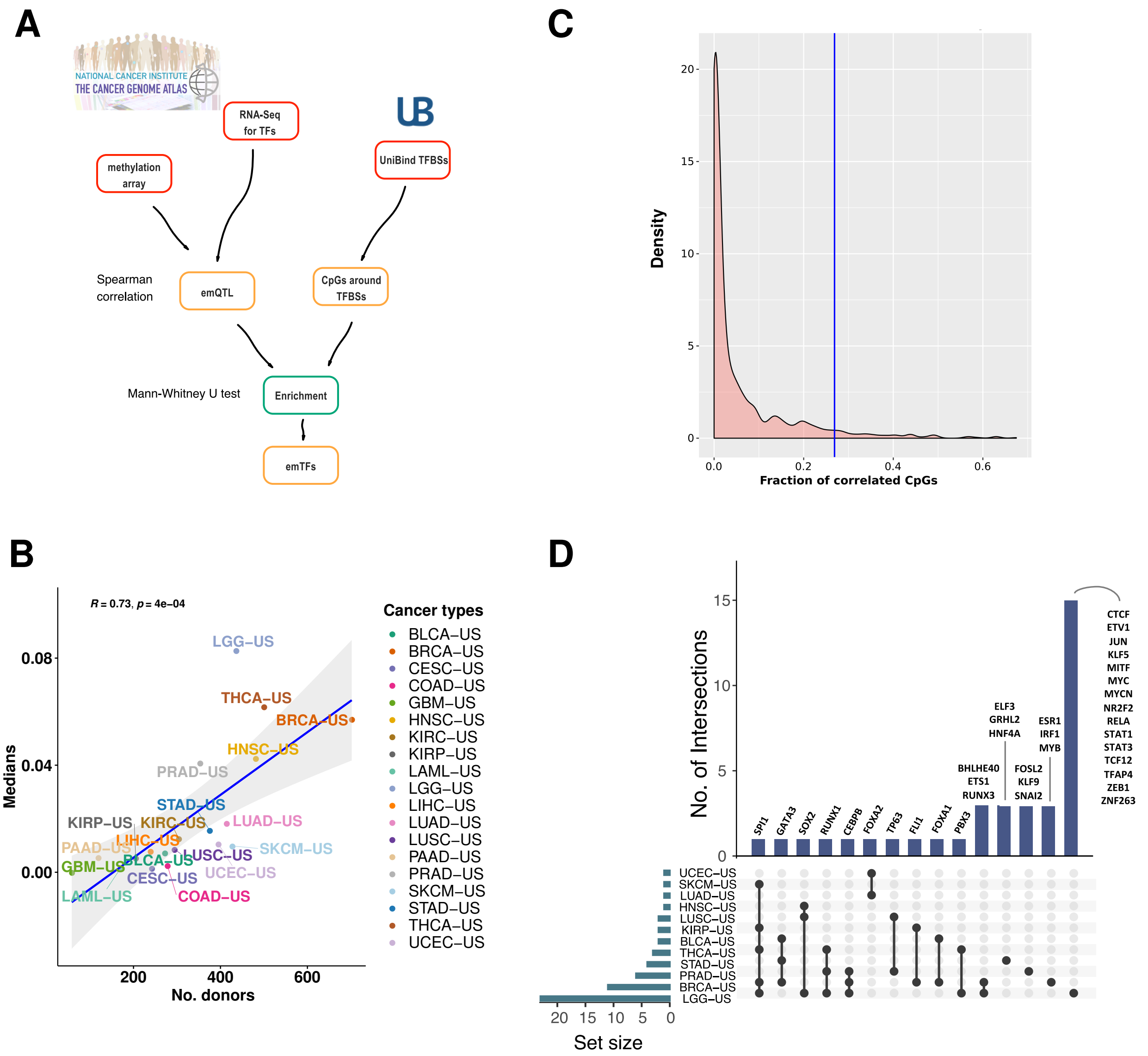


**Figure S1. emQTL analysis identified TFs associated with DNA methylation patterns in TCGA cancer types. A.** Graphical representation of the computational workflow used to carry out the emQTL analysis assessing correlation between DNA methylation at CpGs and TF expression (Materials and Methods). **B.** Density (y-axis) distribution of the proportion of correlated CpGs (x-axis) close to TFBSs from the emQTL analyses when considering all TFs in all cancer types. The vertical blue line represents a threshold at the 95th percentile. **C.** Scatterplot comparing the number of donors/samples per cancer cohort (x-axis) with the median fraction of correlated CpGs from the emQTL analysis per cancer type (y-axis). The blue line represents the fitted Pearson linear relationship with the grey zone representing the 95% confidence interval (Pearson R coefficient and associated p-value are provided in the top-left corner. **D.** Upset plot representing the intersection between the identified TFs in the different cancer types. Each row represents a cancer cohort with points providing information about the intersection (when connected by a vertical line) of the TFs predicted in the different cancer types. The bars at the top indicate the number of intersecting TFs and unique TFs per cancer type. The names of intersecting TFs are annotated above each bar. The set sizes on the left depicts the number of TFs predicted in each cancer type.

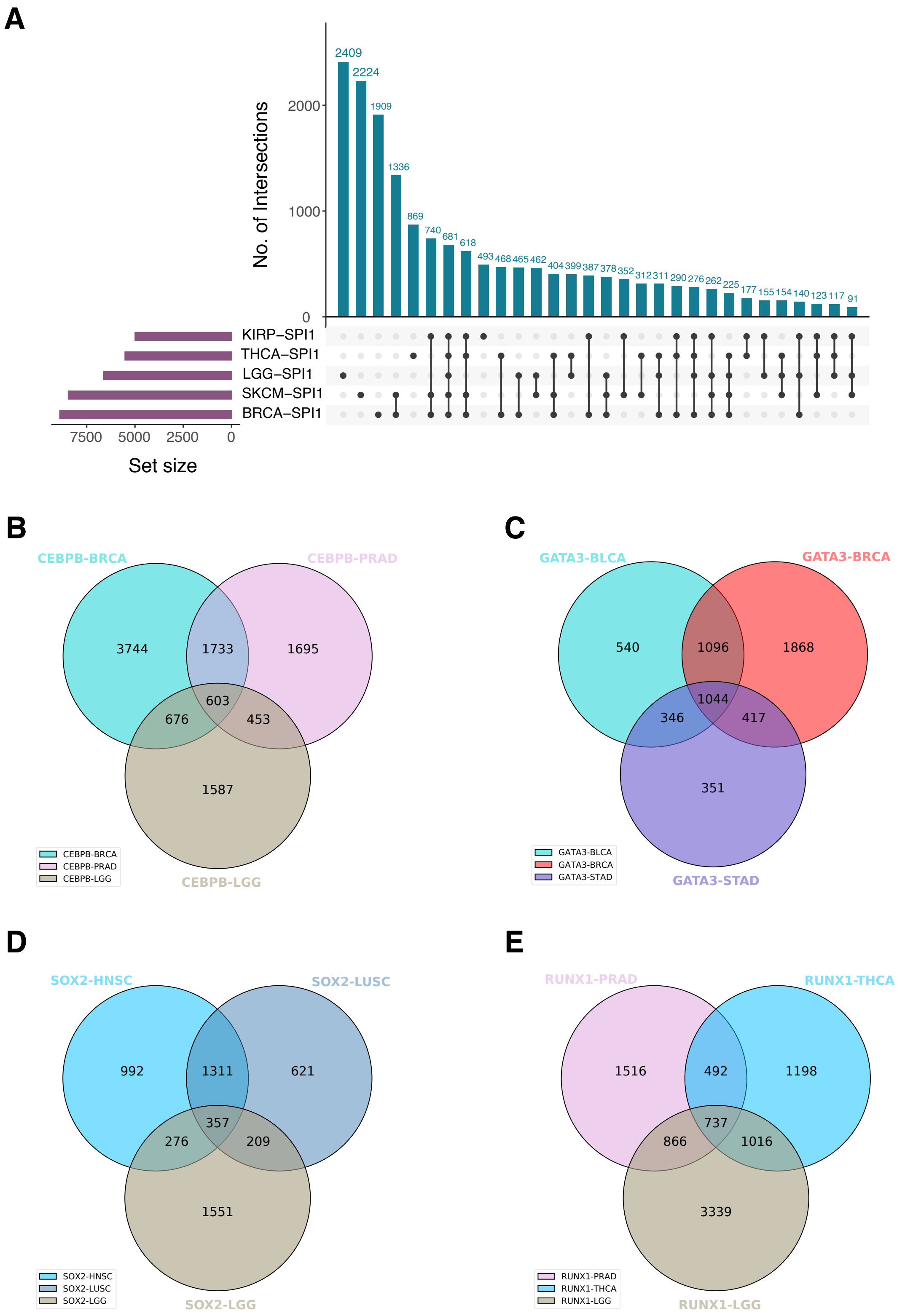


**Figure S2. Assessment of the overlap between emCpGs predicted for the same emTF between cancer types. A.** Upset plot representing the overlap between emCpGs associated with SPI1 in cancer cohorts KIRP, THCA, LGG, SKCM, and BRCA. **B-E** Venn diagrams representing the overlap between emCpGs associated with CEBPB, GATA3, SOX2, and RUNX1, respectively, in the corresponding cancer cohorts (see legend).

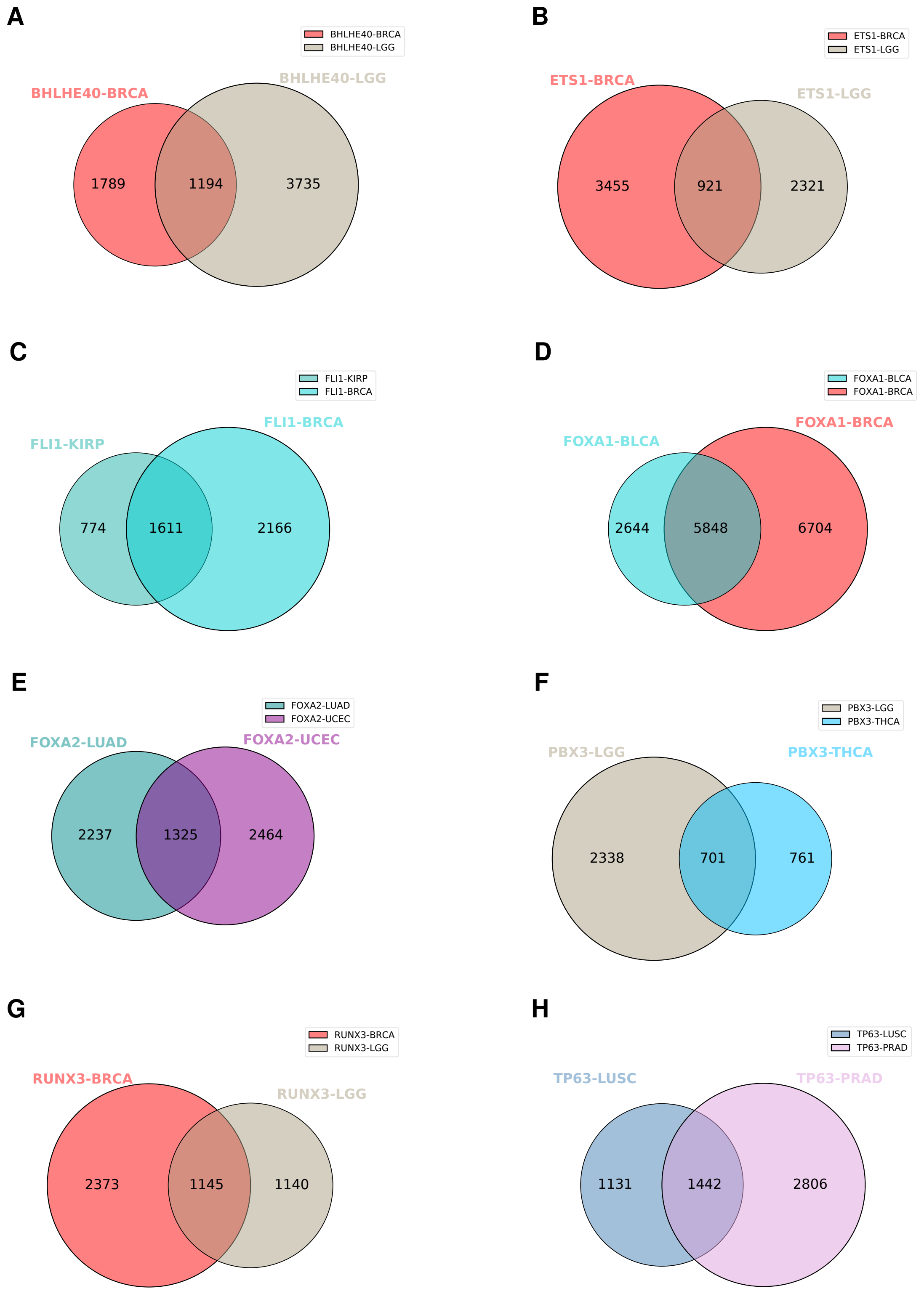


**Figure S3. Assessment of the overlap between emCpGs predicted for the same emTF between cancer types. A-H.** Venn diagrams representing the overlap between emCpGs associated with BHLHE40, ETS1, FLI1, FOXA1, FOXA2, PBX3, RUNX3 and TP63, respectively, in the corresponding cancer cohorts (see legend).

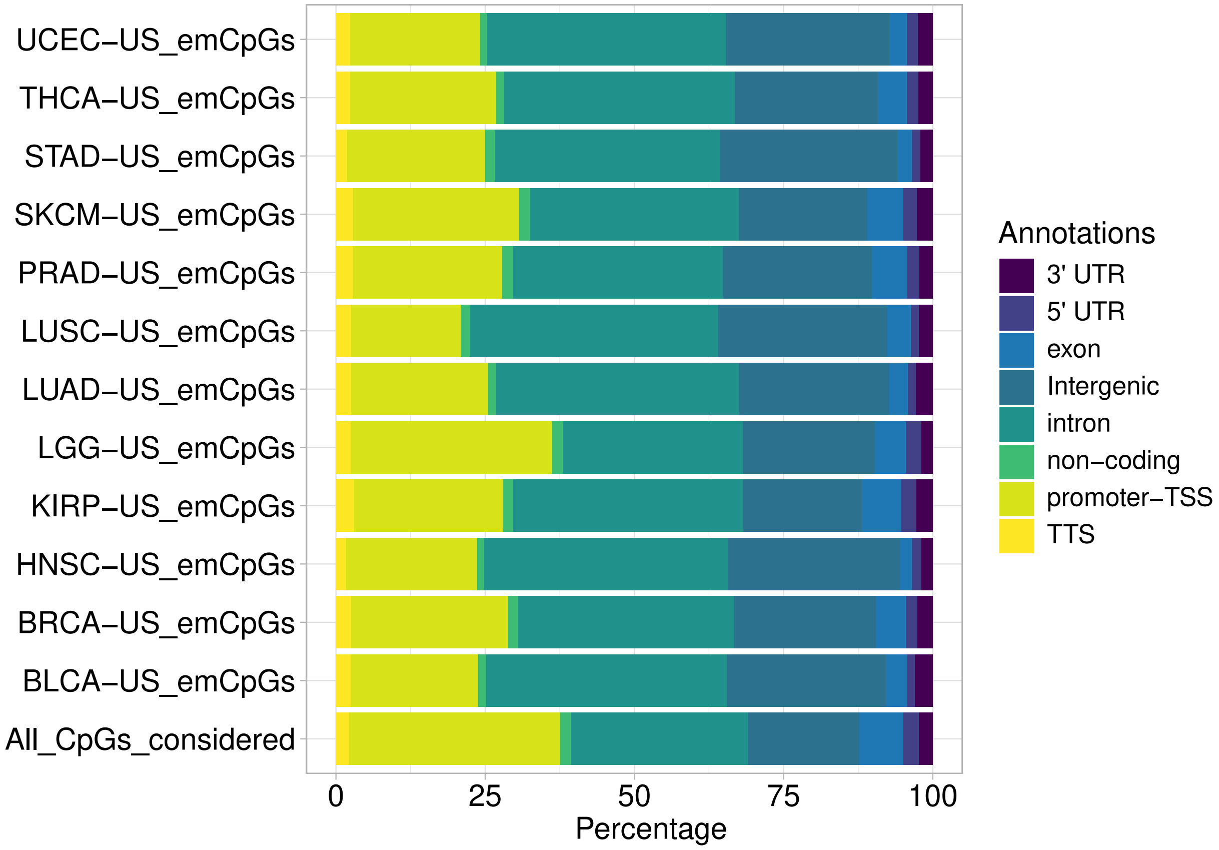


**Figure S4. Functional genomic regions distribution of emCpGs and the complete set of CpGs considered for the emQTL analysis (n=376,997)**. The genomic annotation (see legend) was performed using the *AnnotatePeaks.pl* script from HOMER. The percentage distributions were plotted as horizontal stacked bar plots using the *ggplot2* R package (version 3.3.3).

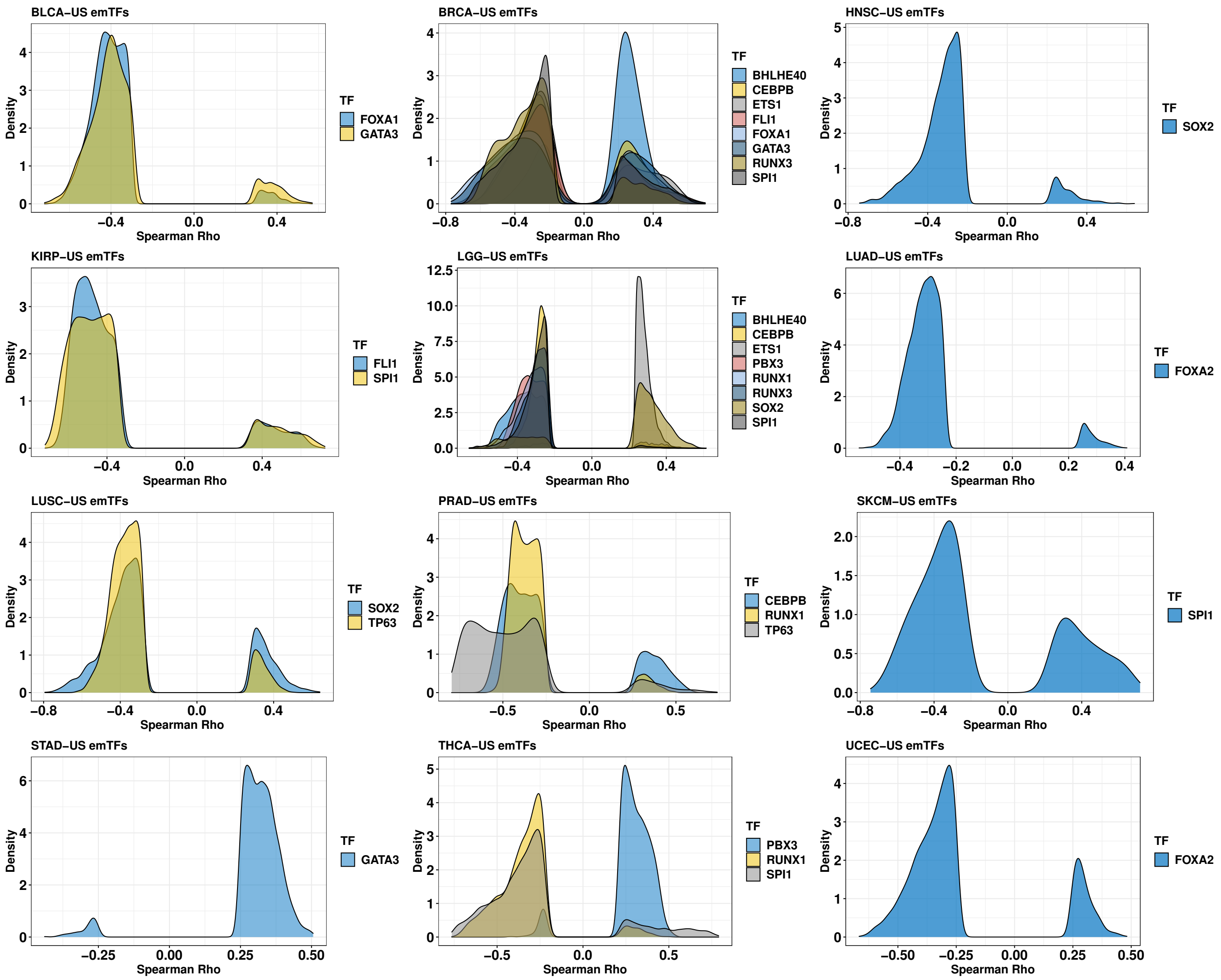


**Figure S5. Distribution of Spearman rho values for emCpGs.** Each panel provides the distribution of the correlation coefficients (Spearman's rho) obtained for the emCpGs associated with the emTFs in different cancer types. Each panel corresponds to the results in a given cancer cohort. When multiple emTFs were identified in a given cancer type, we provide the density plots of each emTF in different colors (see legends).

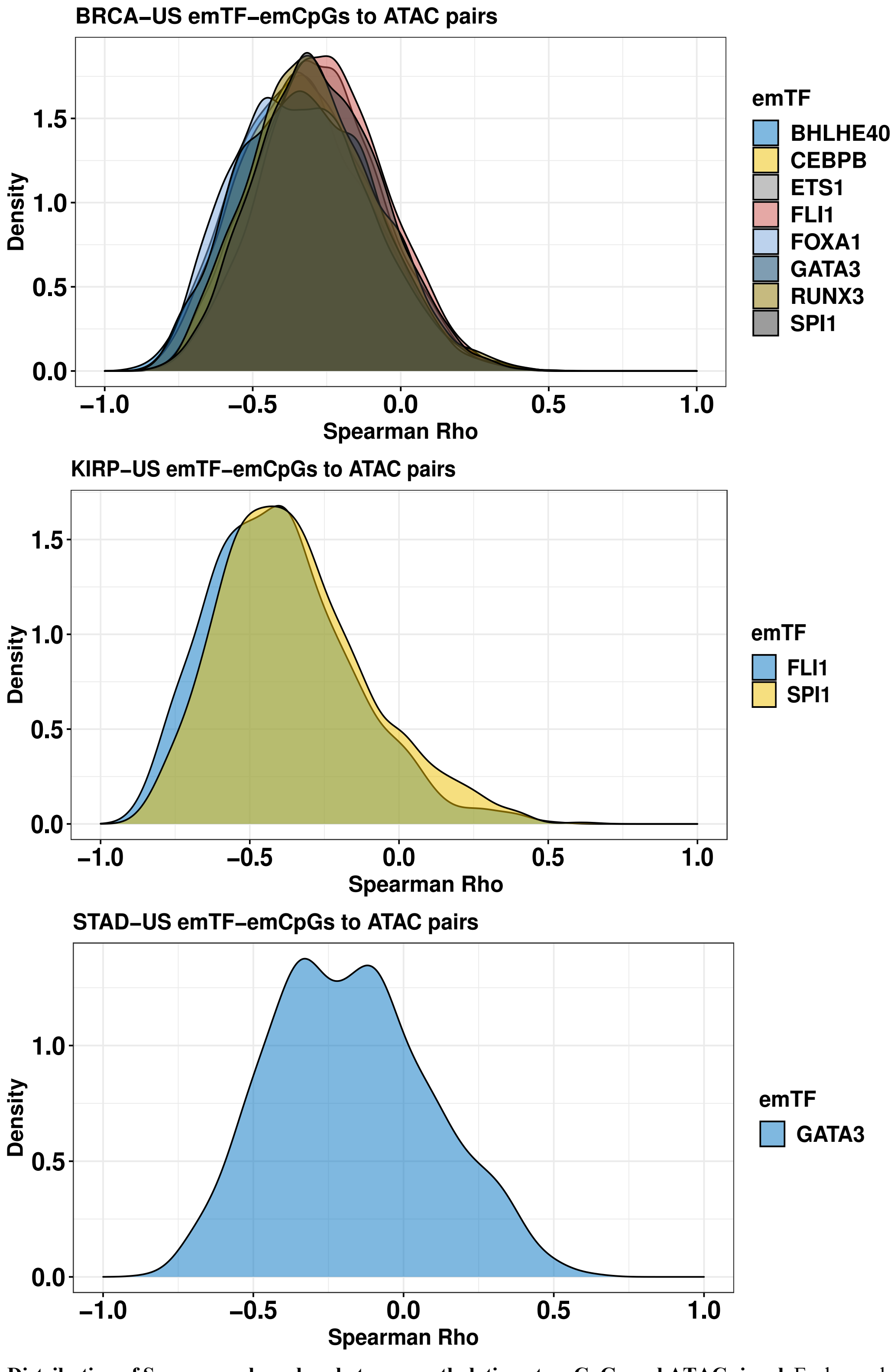


**Figure S6. Distribution of Spearman rho values between methylation at emCpGs and ATAC signal.** Each panel provides the distribution of the pairwise correlation coefficients (Spearman's rho) obtained between emCpGs associated with the emTFs and their paired ATAC regions in three representative cancer types for which at least 20 samples were available. When multiple emTFs were identified in a given cancer type, we provide the density plots of each emTF in different colors (see legends).

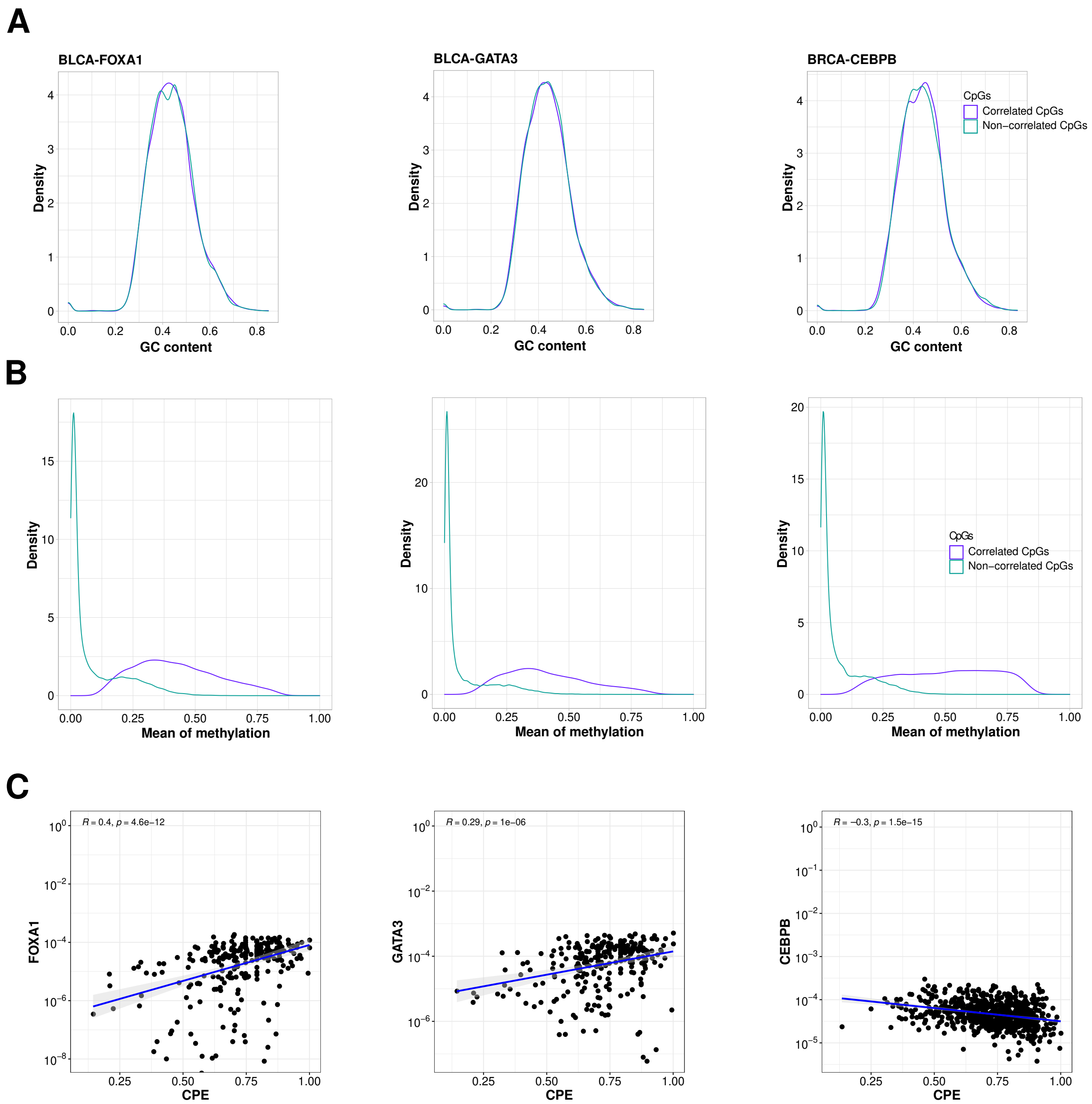


**Figure S7. GC content and DNA methylation profiles for emCpGs and non-correlated CpGs close to the TFBSs of the emTFs. A.** Density distributions of the GC contents computed from regions surrounding emCpGs (purple) and non-correlated CpGs (blue) close to TFBSs of emTFs FOXA1 and GATA3 in the BLCA cohort and CEBPB in the BRCA cohort. **B.** Density distribution of average methylation levels of emCpGs (purple) and non-correlated CpGs (blue) close to FOXA1 and GATA3 in the BLCA cohort and CEBPB in the BRCA cohort**. C.** Scatterplots comparing the tumour purity (x-axis; cumulative purity estimate (CPE) and expression of the TFs (y-axis) for FOXA1, GATA3 in BLCA and CEBPB in BRCA cohorts, respectively. The blue lines represent the fitted Pearson linear relationship with the grey zone representing the 95% confidence interval (Pearson R coefficient and associated p-value are provided in the top-left corner.

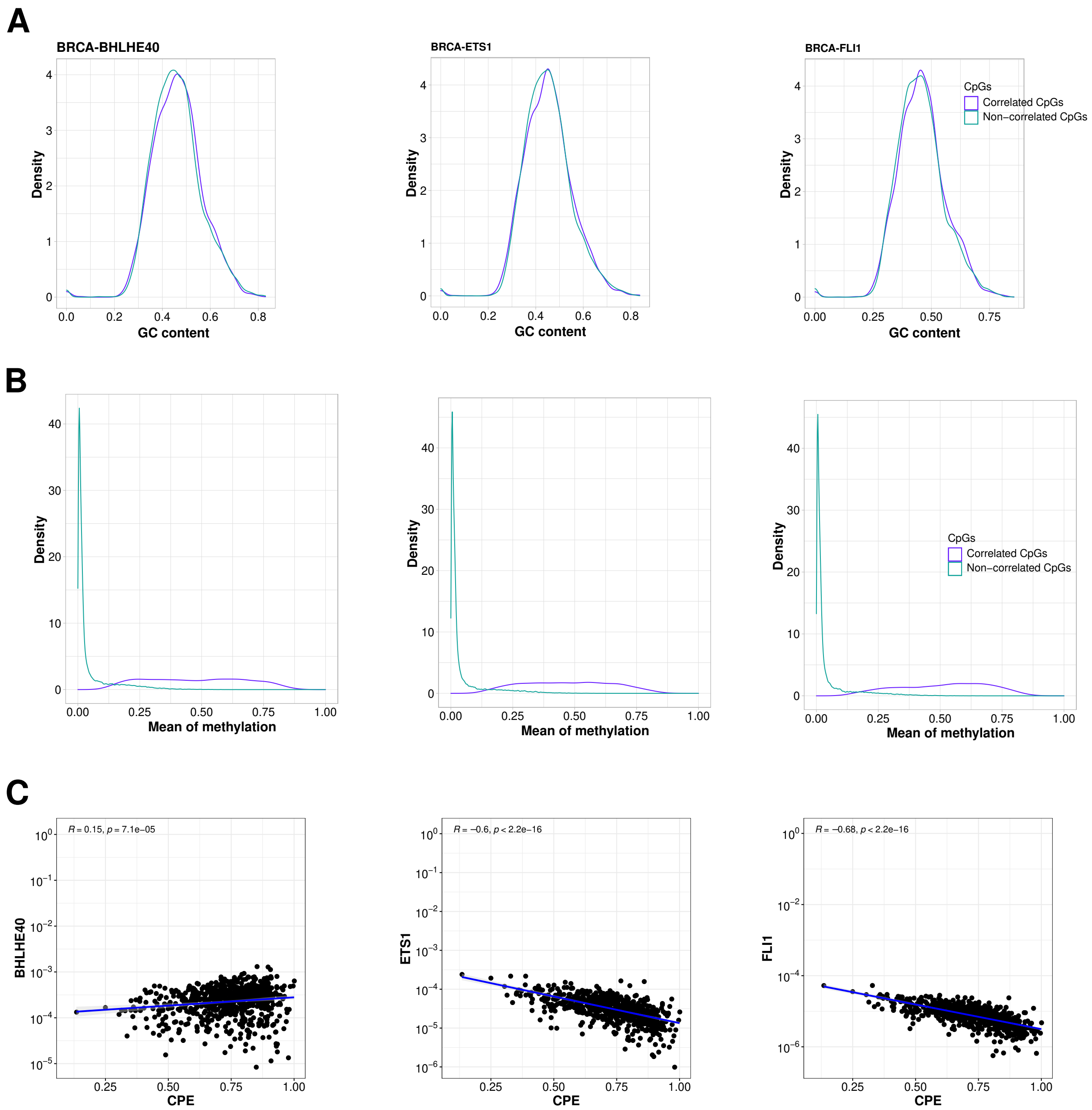


**Figure S8.** Same as Fig. S7 considering BHLHE40, ETS1, and FLI1 in the BRCA cohort.

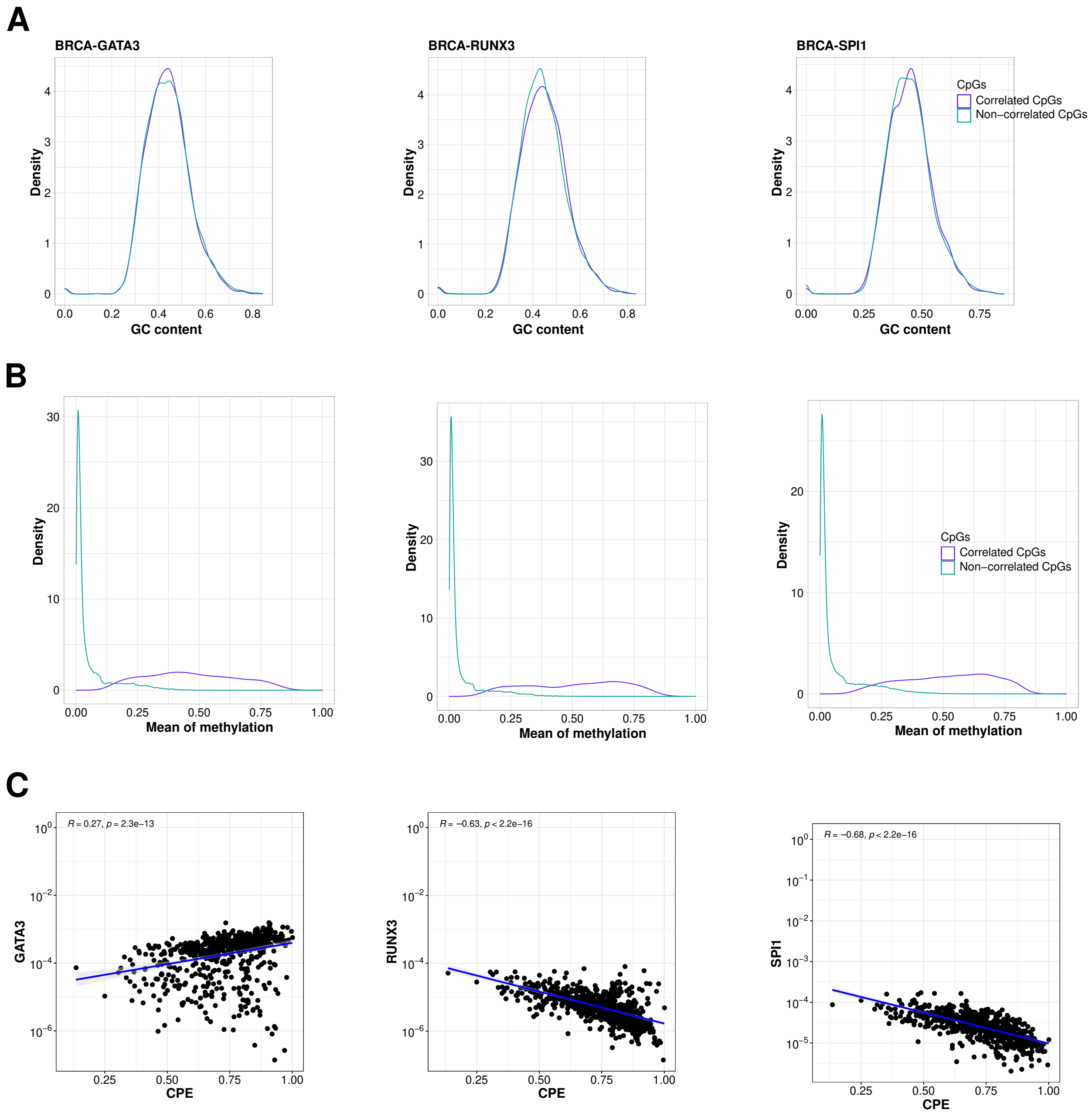


**Figure S9.** Same as Fig. S7 considering GATA3, RUNX3, and SPI1 in the BRCA cohort.

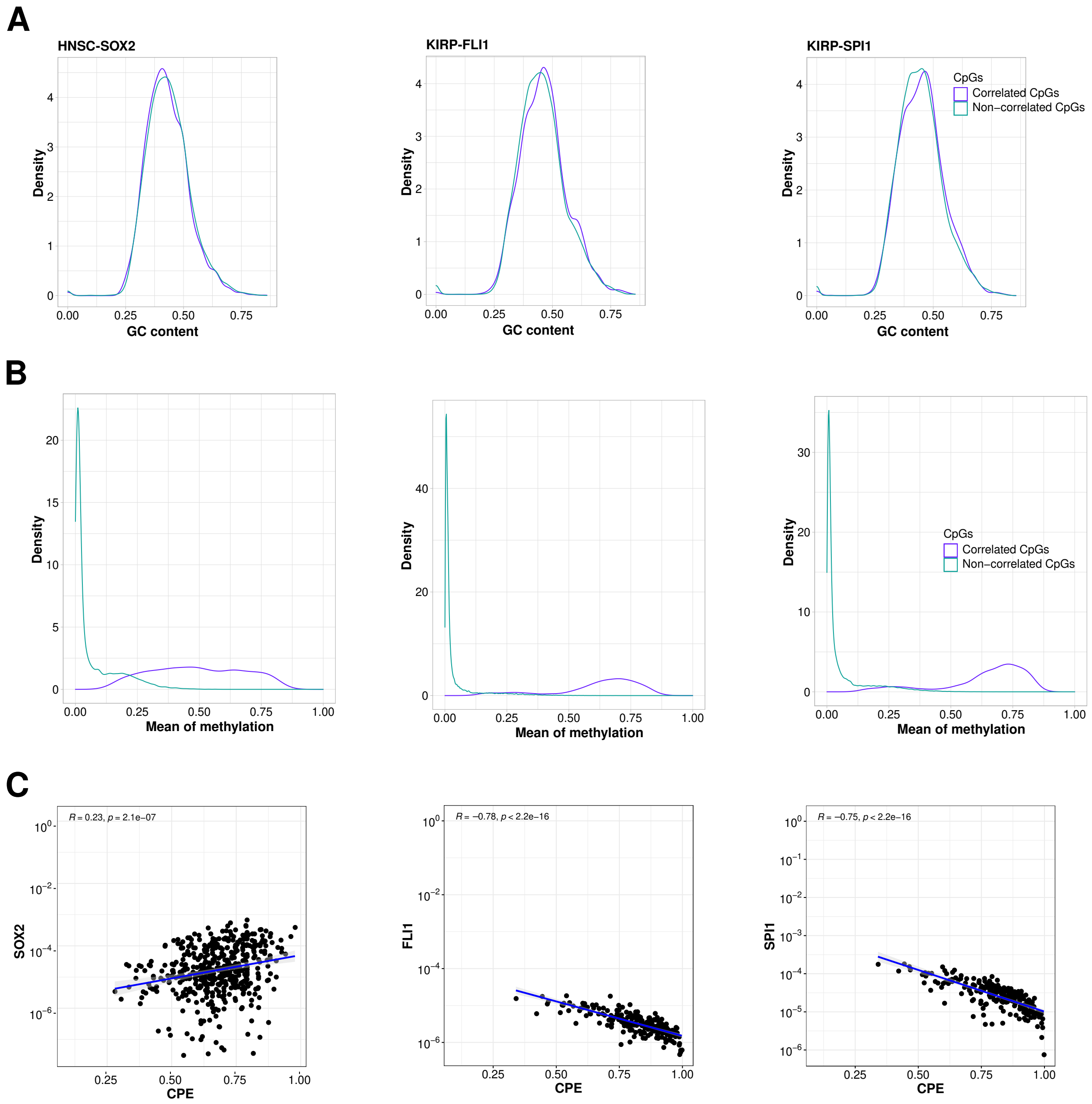


**Figure S10.** Same as Fig. S7 considering SOX2 in the HNSC cohort, and FLI1 and SPI1 in the KIRP cohort.

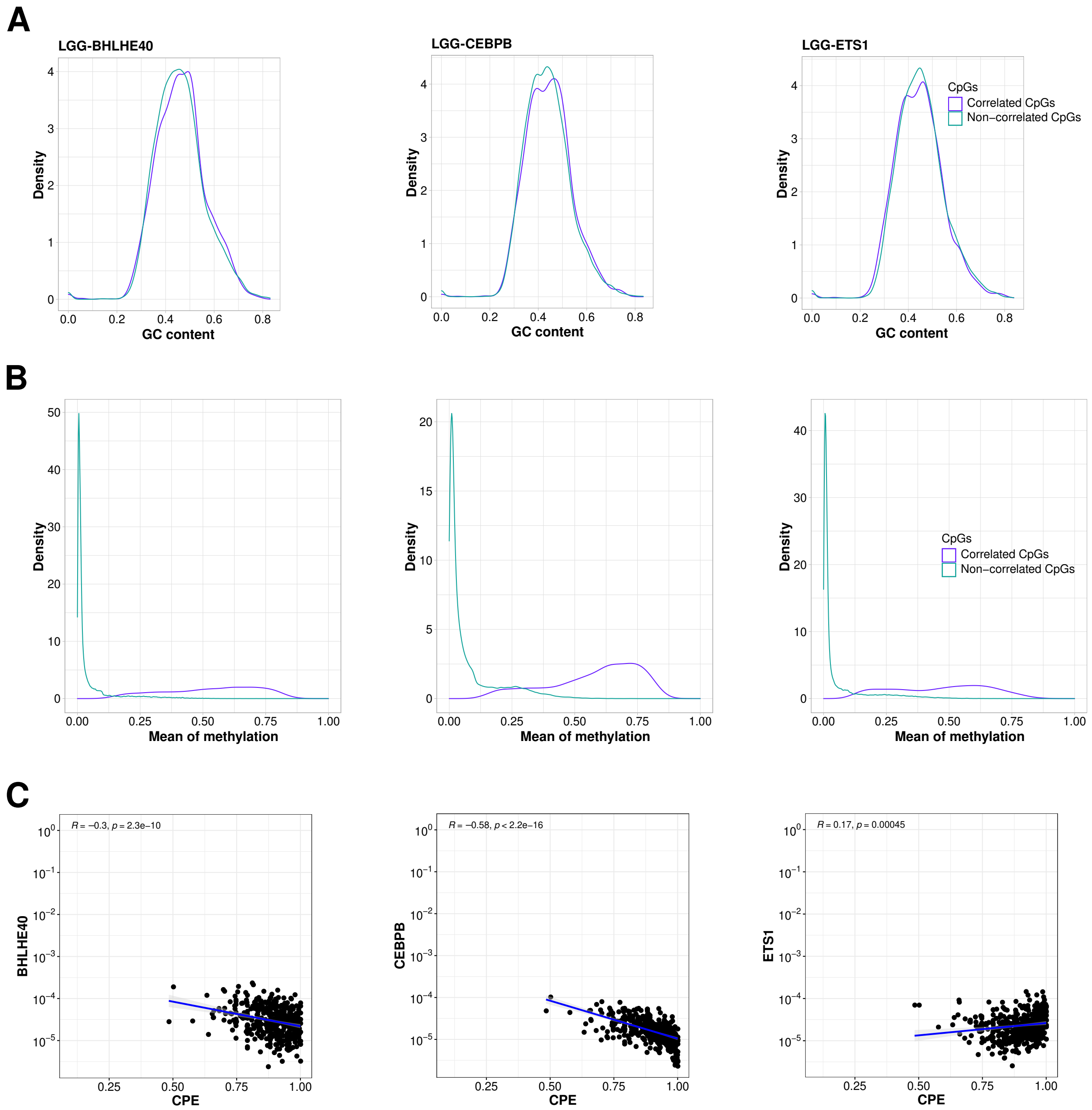


**Figure S11.** Same as Fig. S7 considering BHLHE40, CEBPB, and ETS1 in the LGG cohort.

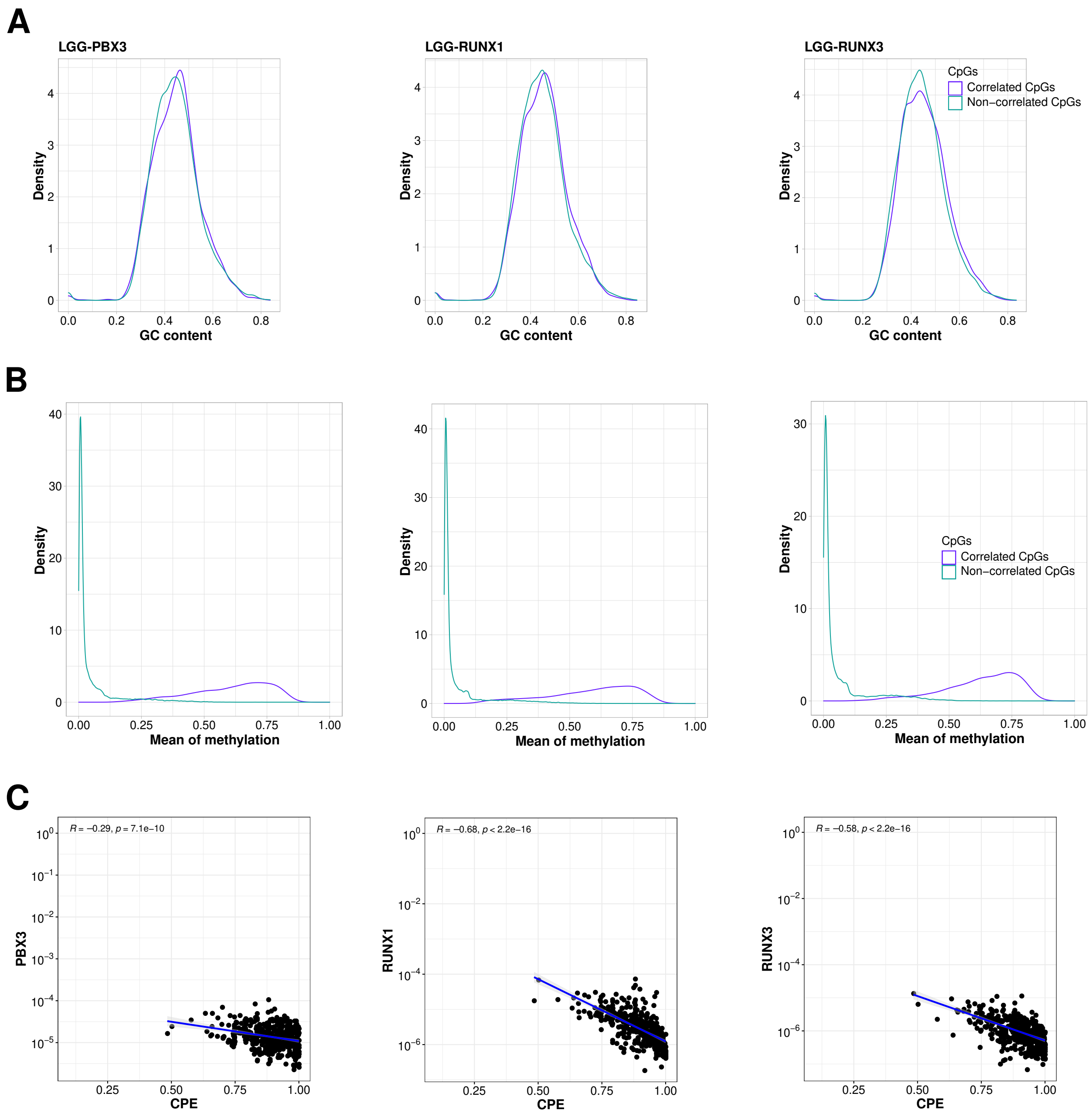


**Figure S12.** Same as Fig. S7 considering PBX3, RUNX1, and RUNX3 in the LGG cohort.

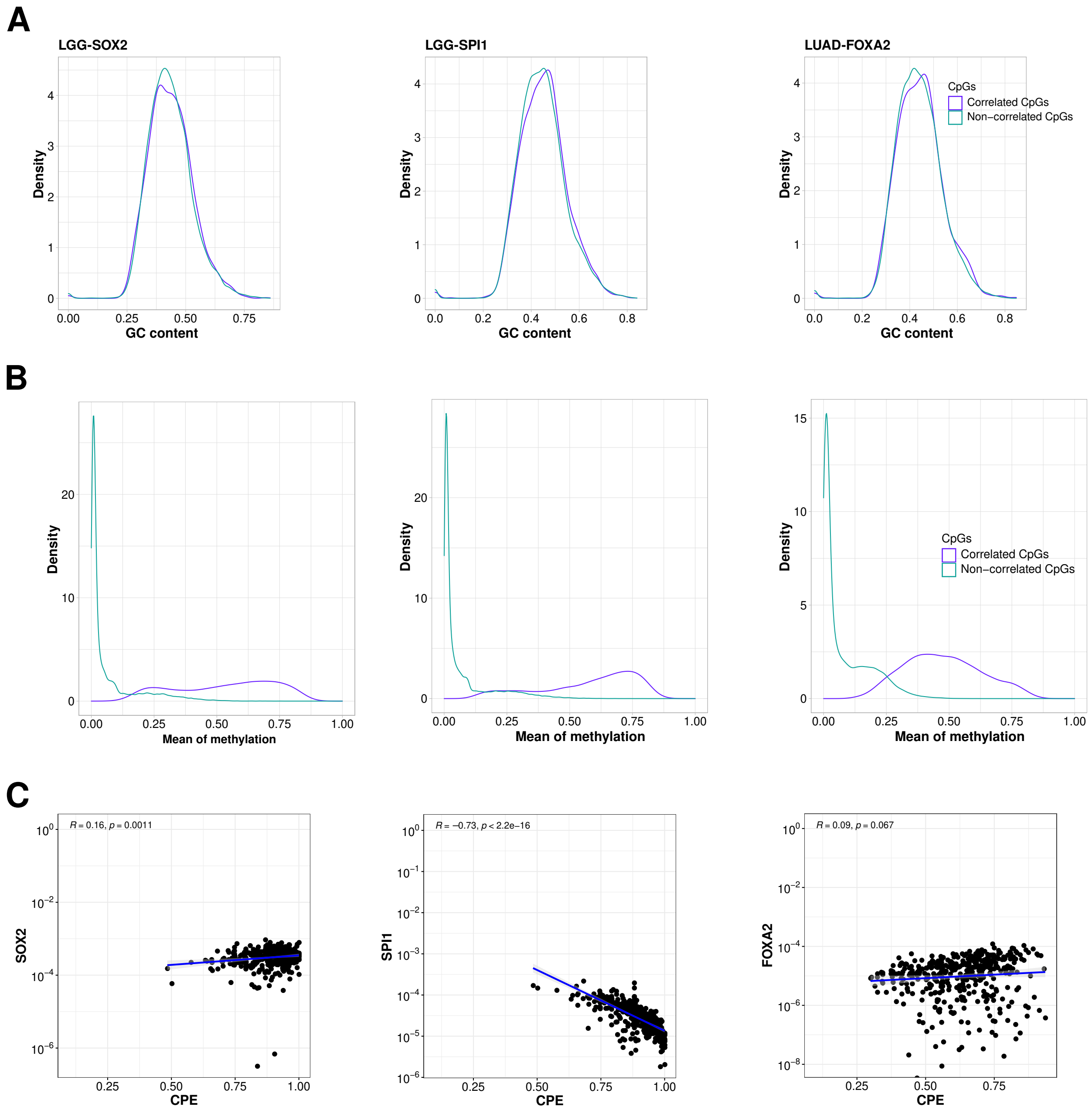


**Figure S13.** Same as Fig. S7 considering SOX2 and SPI1 in the LGG cohort and FOXA2 in the LUAD cohort.

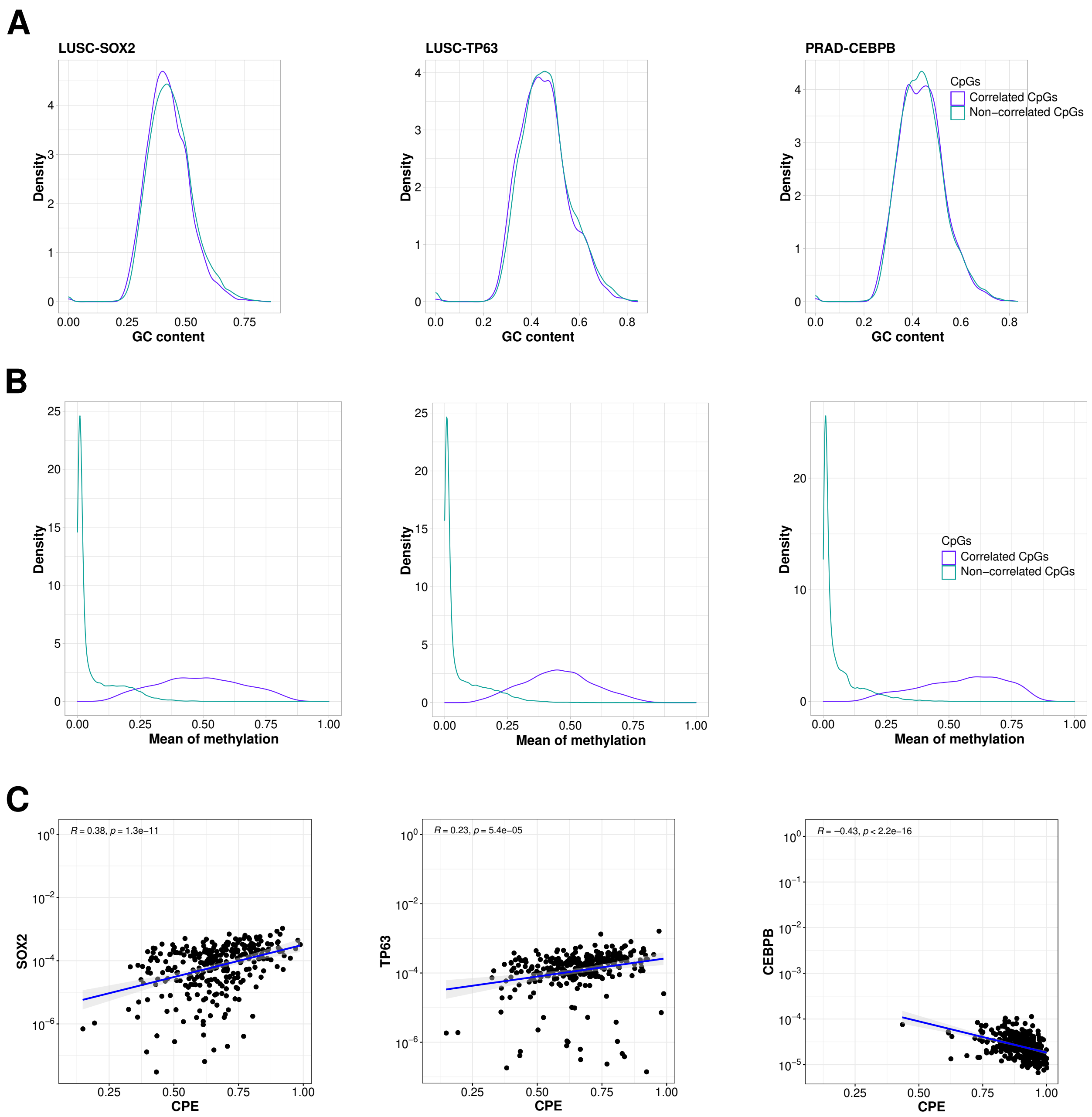


**Figure S14.** Same as Fig. S7 considering SOX2 and TP63 in the LUSC cohort, and CEBPB in the PRAD cohort.

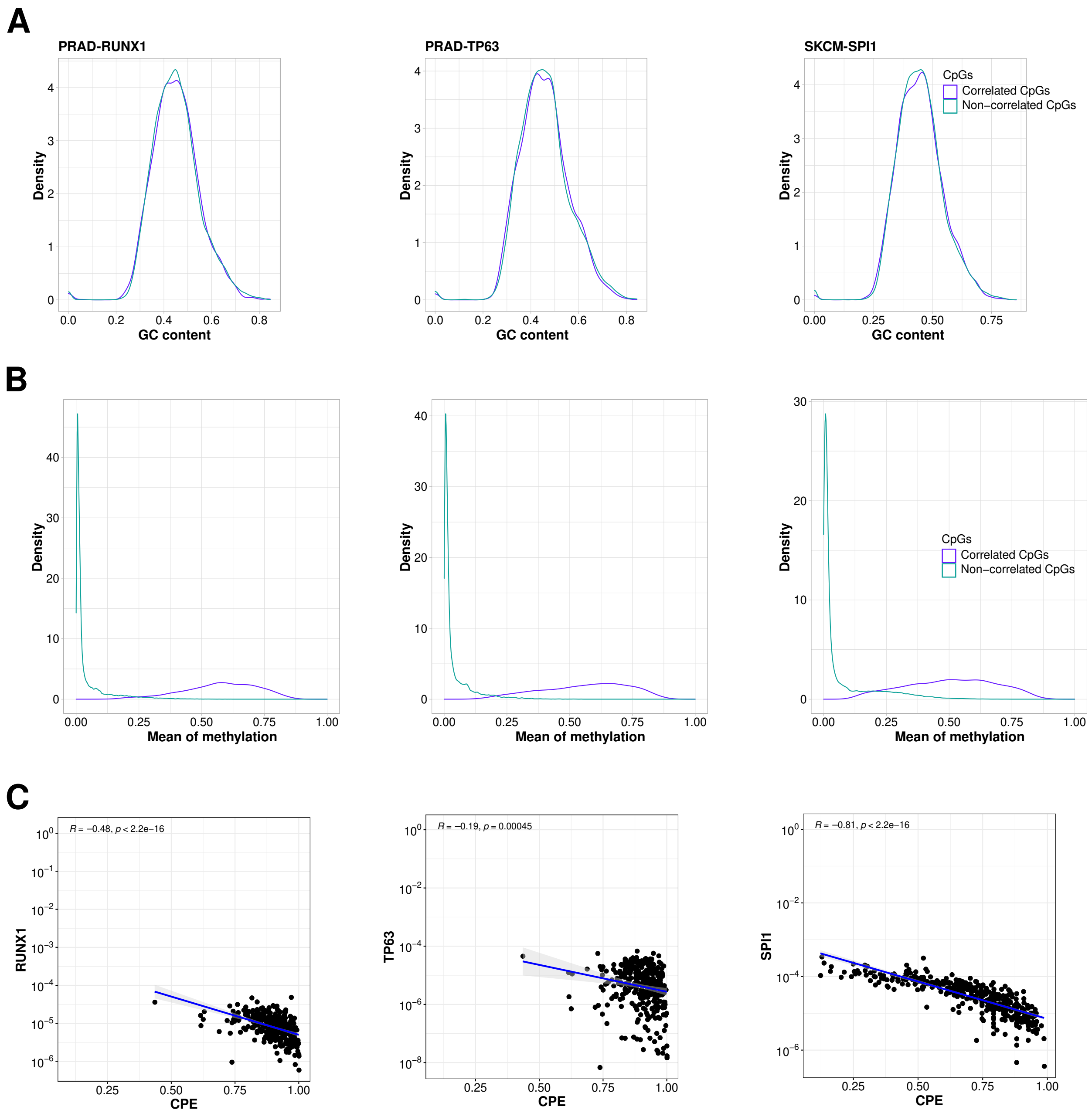


**Figure S15.** Same as Fig. S7 considering RUNX1 and TP63 in the PRAD cohort, and SPI1 in the SKCM cohort.

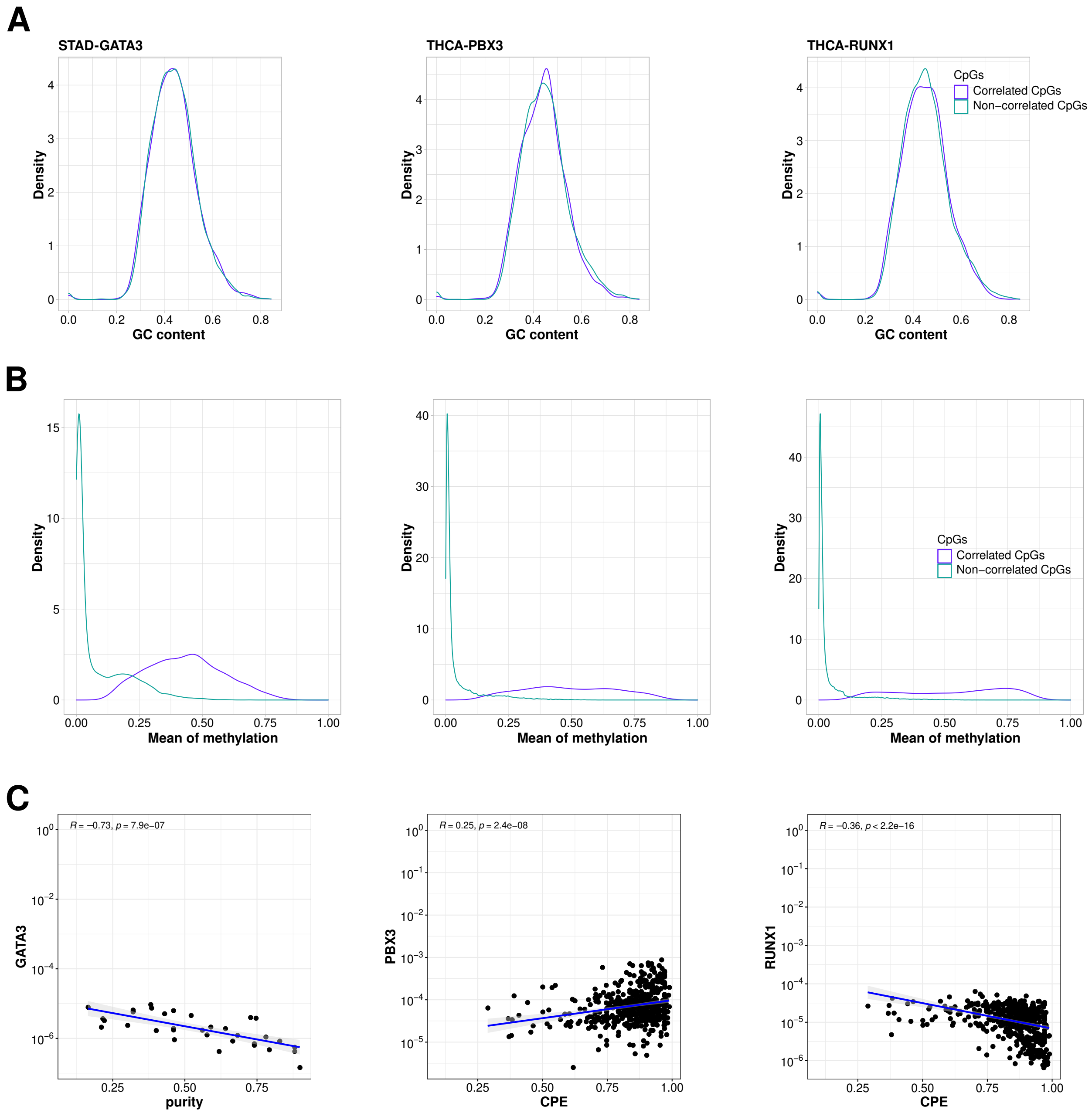


**Figure S16.** Same as Fig. S7 considering GATA3 in the STAD cohort, and PBX3 and RUNX1 in the THCA cohort.

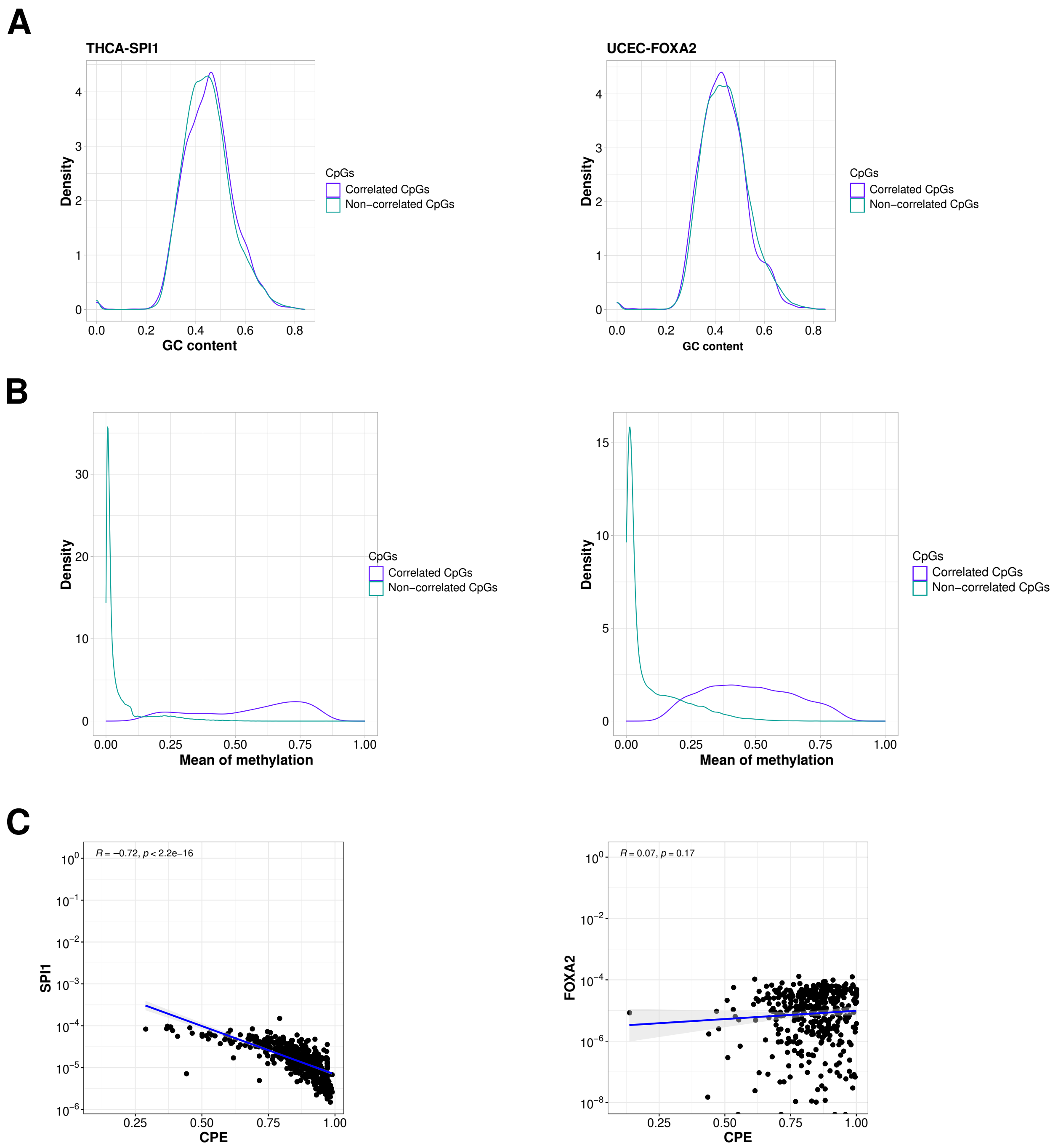


**Figure S17.** Same as Fig. S7 considering SPI1 in the THCA cohort and FOXA2 in the UCEC cohort.

**A**

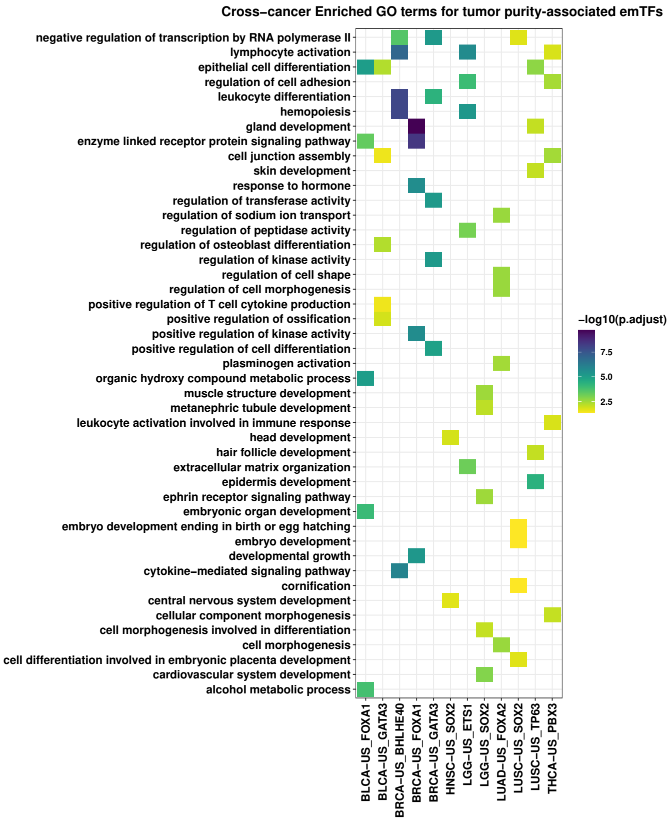


**B**

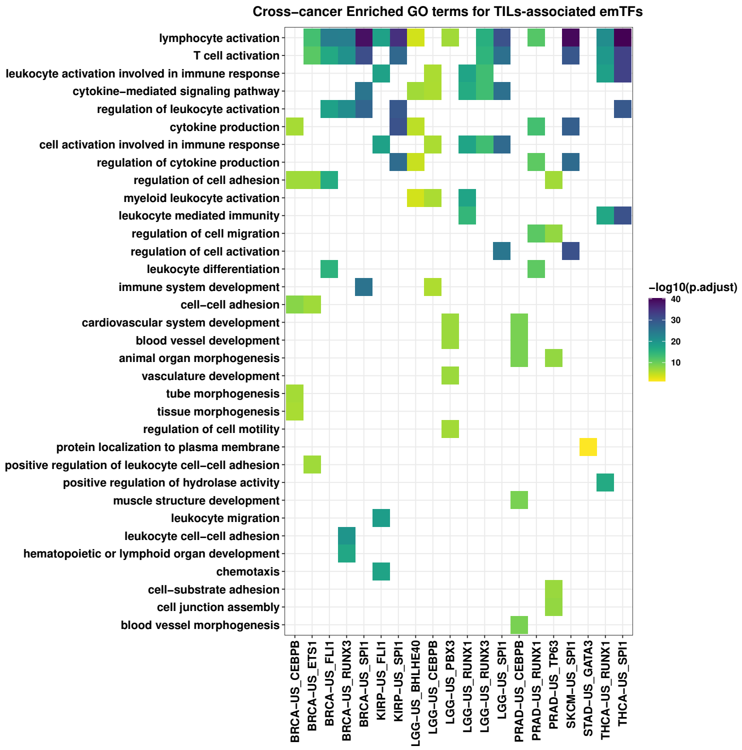


**Supplementary Figure S18. GO enrichment for gene targets of emCpGs. A.** Top 5 enriched GO biological process terms (ranked by adjusted p-values) for each emTF-cancer pair are represented when considering target genes of CpGs associated with emTFs whose expression showed positive correlation with tumor purity. **B.** Same as in A. when considering emTFs whose expression showed negative correlation with tumor purity.

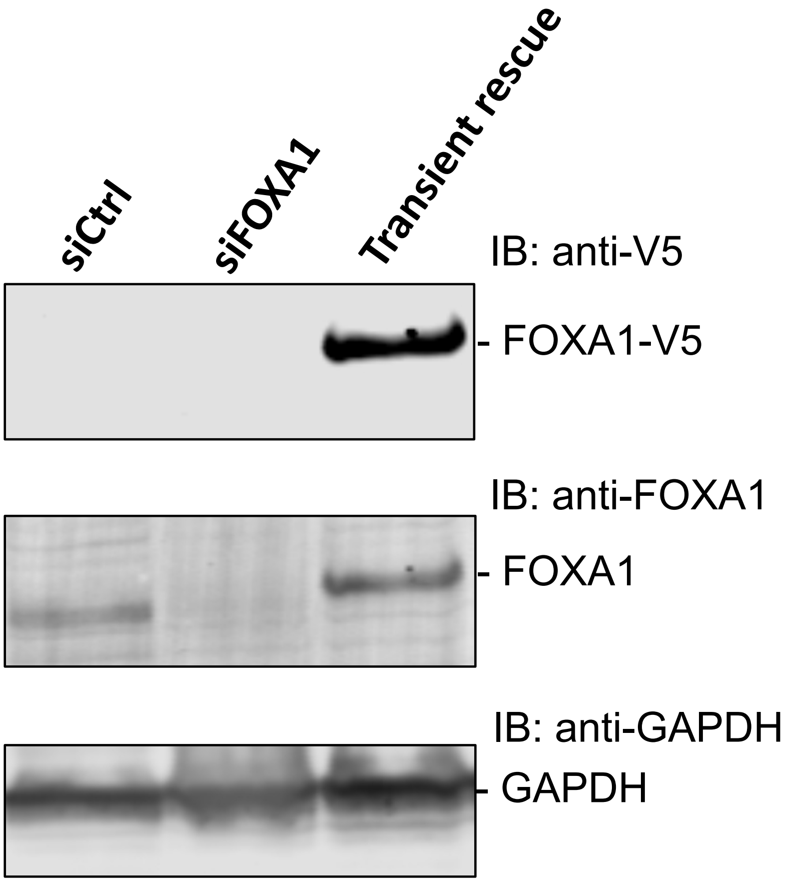


**Figure S19. Western blot validations of siFOXA1 and transient rescue.** As a negative control, we used siCtrl (AllStars Negative Control siRNA from Qiagen) (first column). The western blot of the siRNA-mediated knockdown (KD) of endogenous FOXA1 expression in MCF-7 cells is provided in the second column. The western blot of the siFOXA1-mediated KD rescued with transient expression of FOXA1-V5 is provided in the third column. We used rabbit anti-FOXA1 polyclonal, mouse anti-V5 monoclonal, and mouse anti-GAPDH polyclonal antibodies. GAPDH was probed as loading control.

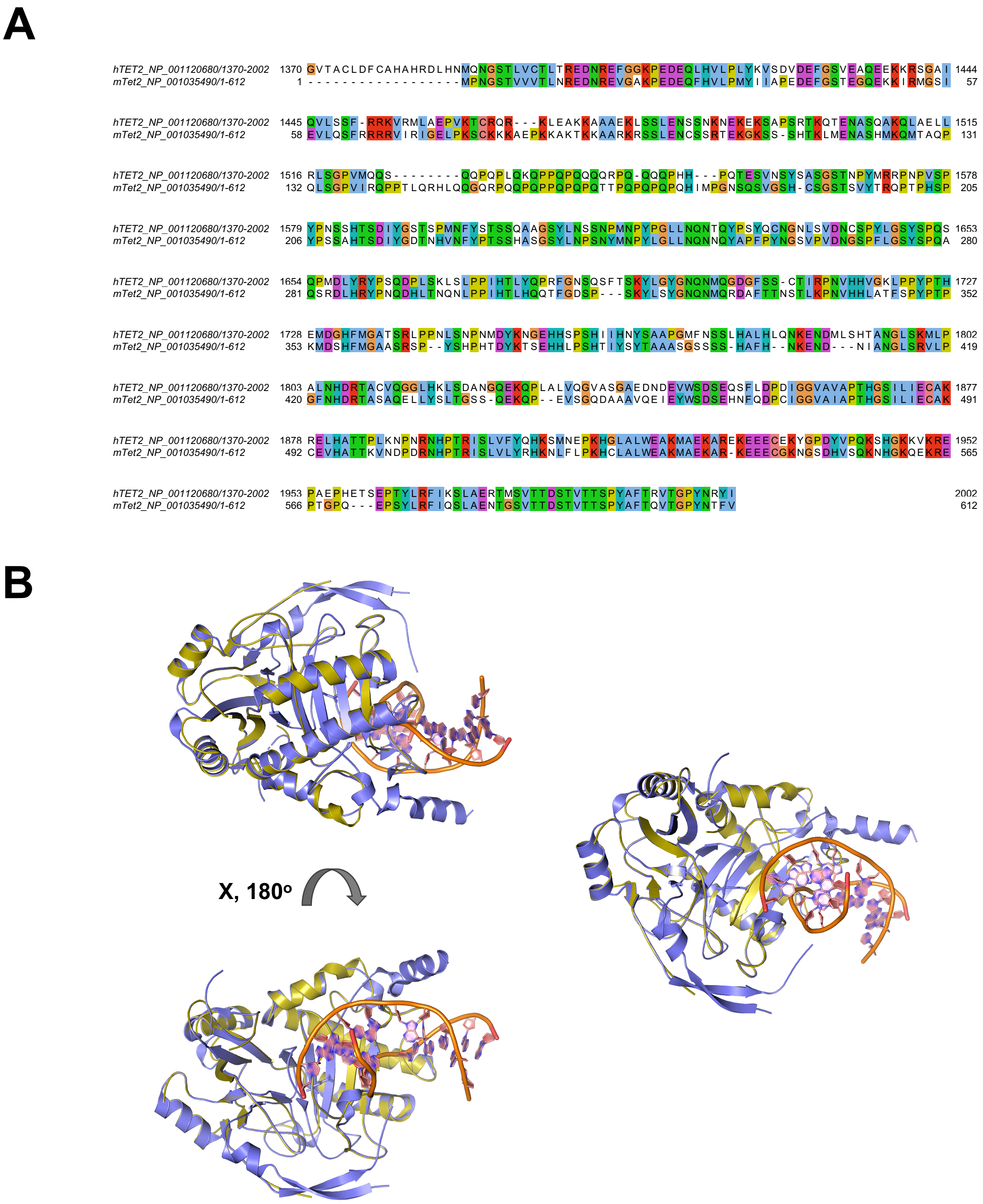


**Figure S20. Comparison between mTET2 and hTET2. A.** Pairwise sequence alignment of mTET2 isoform 2 (RefSeq ID: NP_001035490) and hTET2 isoform1 (RefSeq ID: NP_001120680). The pairwise sequence alignment of the two proteins was investigated using the MUSCLE algorithm accessed through Jalview. We observe a pairwise sequence identity of 59.72 %. **B.** Structural alignment of hTET2 (PDB ID: 4nm6A), cartoon rendering in slate blue, and modelled structure of mTET2 (PDB ID: Q6NO21), cartoon rendering in yellow. The two structures superimpose well on top of each other, indicating that these two proteins are structurally similar. Cartoon rendering of the structures was performed using pyMOL version 2.1.
